# Comprehensive maps of escape mutations from antibodies 10-1074 and 3BNC117 for Envs from two divergent HIV strains

**DOI:** 10.1101/2025.01.30.635715

**Authors:** Caelan E. Radford, Jesse D. Bloom

## Abstract

Antibodies capable of neutralizing many strains of HIV are being explored as prophylactic and therapeutic agents, but viral escape mutations pose a major challenge. Efforts have been made to experimentally define the escape mutations from specific antibodies in specific viral strains, but it remains unclear how much the effects of mutations on neutralization differ among HIV strains. Here, we use pseudovirus deep mutational scanning to comprehensively map escape mutations from the V3 loop targeting antibody 10-1074 and the CD4-binding site targeting antibody 3BNC117 for both a clade A (BF520) and a clade B (TRO.11) HIV Envelope (Env). Mutations that escape neutralization by antibody 10-1074 are largely similar for the two Envs, but mutations that escape 3BNC117 differ greatly between Envs. Some differences in the effects of mutations on escape between Envs can be explained by strain-to-strain variation in mutational tolerance or glycosylation patterns, but other mutations have different effects on escape for unclear reasons. Overall, the extent that measurements of mutational effects on antibody neutralization can be generalized across HIV strains differs among antibodies.

**Importance:** Broadly neutralizing antibodies are promising candidates as prophylactics and therapeutics for HIV. This study uses pseudoviruses to map all escape mutations for antibodies 10-1074 and 3BNC117 for the Envelope proteins from two different HIV strains. These maps can inform analyses of viral mutations observed in clinical trials, and help understand how the escape mutations from these antibodies differ across HIV strains.

## Introduction

Antibodies capable of broadly neutralizing diverse strains of HIV are being explored as prophylactic and therapeutic treatments, and have shown promise in suppressing viremia or preventing transmission of antibody-sensitive viruses^1,2^. V3 loop targeting antibody 10-1074^3^ and CD4-binding site targeting antibody 3BNC117^4^ have been used in singlet and in combination for clinical trials to suppress viremia in individuals living with HIV^5–10^. Antibody 10-1074 neutralization of HIV Env is dependent on a glycosylation at site N332 and antibody-Env interactions at a co-receptor binding site spanning residues 324 to 327^3,11^.

Antibody 3BNC117 neutralizes HIV Env through interactions at residues dispersed across the CD4-binding site at loop D, the CD4-binding loop, and near V5, as well as at site 207 and sites in the V3 loop of the adjacent protomer^12–14^. Due to their non-overlapping epitopes, the combination of 10-1074 and 3BNC117 for treatments makes it less likely for viruses to acquire resistance through single mutations.

Even when structures of antibodies bound to Env are available, it can be challenging to predict which Env mutations will escape antibody neutralization. Mutations that cause escape from neutralization can be outside of the structurally defined antibody epitope^14–16^, some mutations at sites within the structurally defined epitope may not affect neutralization^14–18^, and some mutations in the epitope may not be tolerable for Env folding or function. Furthermore, the effects of mutations on antibody neutralization can vary among HIV strains, which can have substantially divergent Env proteins^19^. These strain-to-strain differences can be due to differing mutational tolerance between Envs^20^, differences in Env glycosylation or variable loop length^21^, or other differences in Env conformation or dynamics^22,23^.

Deep mutational scanning is an experimental method that can be used to measure the effects of all mutations to a viral entry protein on its cell entry function of neutralization by an antibody^18,24,25^. We previously developed a lentivirus deep mutational scanning platform that enables measurement of the effects of all mutations to HIV Env using pseudoviruses that can only undergo a single round of cellular infection and so provide a safe tool to study viral protein mutations at biosafety level 2^18^. We have previously used this platform to measure how mutations to the Env of the clade A transmitted/founder HIV strain BF520.W14M.C2 Env (hereafter referred to as BF520) affect its cell entry function and neutralization by a set of CD4-binding site targeting antibodies, including antibody 3BNC117^26^. Here we use the deep mutational scanning platform to measure the effects of all mutations to the Env of the clade B HIV strain TRO.11 Env on cell entry and neutralization by antibodies 10-1074 and 3BNC117, as well as the effects of all mutations to the BF520 Env on neutralization by 10-1074. We then analyze these data to compare the escape mutations for the two antibodies across the two Envs to assess the extent to which escape mutations from the same antibody are similar or different for Envs from divergent strains.

## Results

### Choice of two strains of HIV Env from different clades

We selected Envs from two divergent strains of HIV for deep mutational scanning: clade A BF520 and clade B TRO.11 (Figure 1A)^27–29^. These Envs have 73% percent amino acid identity, and differ by 237 amino-acid substitutions, insertions, and deletions (Figure 1A,B).

**Figure 1:**
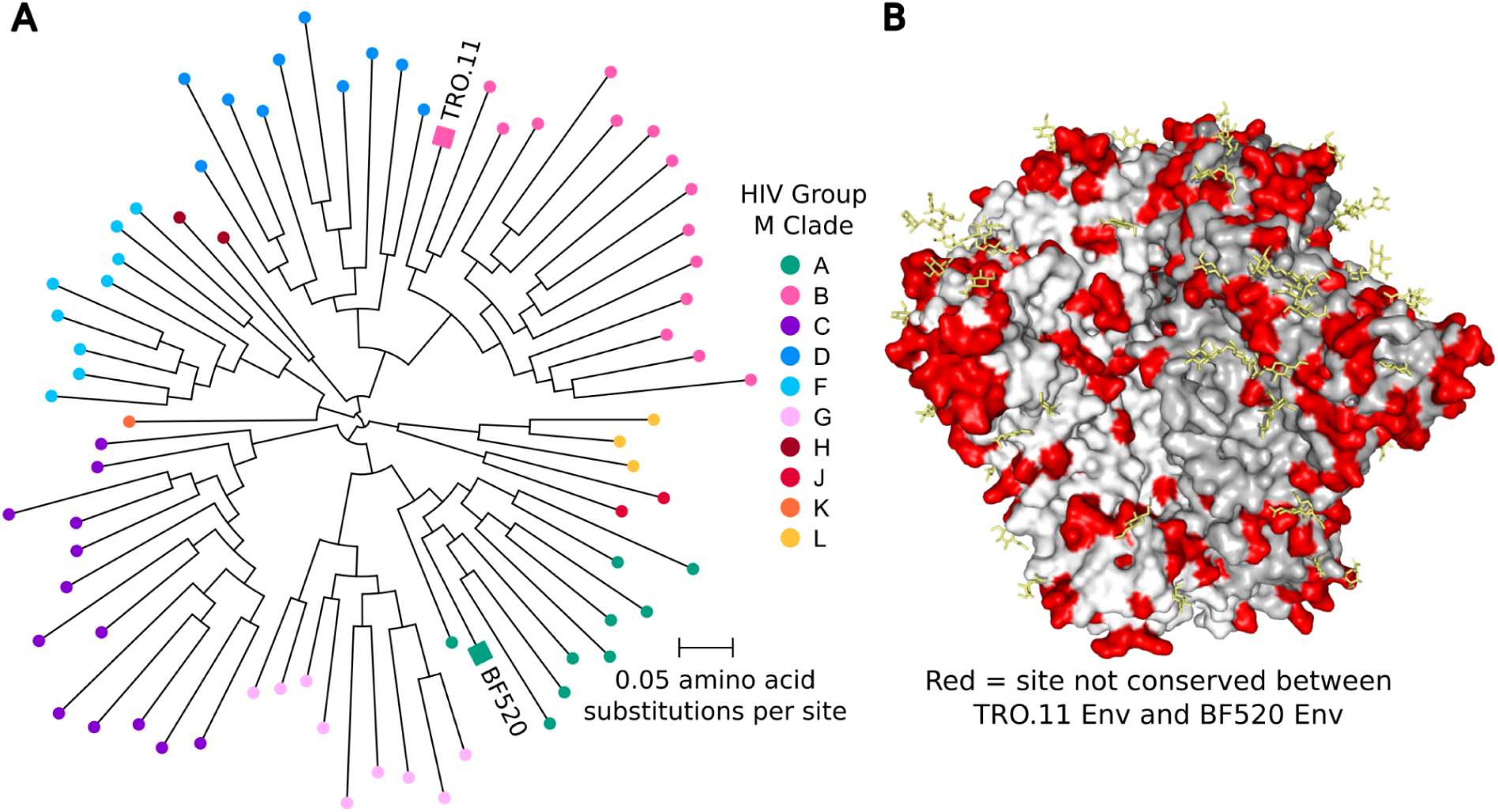
Divergence of Env strains TRO.11 and BF520. (A) HIV group M unrooted phylogenetic tree highlighting strains TRO.11 and BF520. The tree was inferred using all Env sequences from the 2022 HIV Sequence Compendium alignment of HIV group M Env protein sequences from the Los Alamos National Laboratory HIV Sequence Database^39^. The phylogenetic tree was inferred using IQ-TREE^40^ with the HIV-B_m_ amino acid substitution model^41^, and rendered using ETE3^42^. (B) Structure of HIV Env (9BER^43^) with sites that differ in their amino-acid identity between TRO.11 Env and BF520 Env colored red. Individual gp120-gp41 heterodimers of Env are shaded white or gray for clarity. Glycans that are present in the structure are colored yellow.

We have previously performed deep mutational scanning of the transmitted-founder clade A BF520 Env^26^, and the effects of BF520 Env mutations on cell entry and neutralization by antibody 3BNC117^4^ analyzed here are from that prior study^26^. The previously described BF520 Env mutant libraries from that prior study were used in new experiments in this paper to measure the effects of mutations on escape from antibody 10-1074.

To generate data on the effects of mutations to a clade B Env, we performed entirely new deep mutational scanning for this study. We selected TRO.11 Env as a representative clade B Env for these new experiments because it is on a virus panel widely used to assess anti-HIV antibody neutralization breadth^30^ and yields relatively high virus titer in the pseudovirus system used in our study^26^.

### TRO.11 Env deep mutational scanning mutant library

The TRO.11 deep mutational scanning was performed using the same previously described pseudovirus system^18^ that we recently applied to BF520 Env^26^. The system enables the generation of single-round infectious lentivirus mutant libraries where each barcoded variant in a library has a genotype-phenotype link between a nucleotide barcode in its genome and the Env mutant on its surface^18,26^.

We made two replicate site-saturation mutant libraries covering the TRO.11 Env ectodomain (Supplemental Figure 1A-C). Together, these libraries contained 183,593 barcoded variants and contained 97% of all possible amino-acid mutations to the TRO.11 Env ectodomain (Supplemental Figure 1A). The Envs in the libraries had an average of one mutation per barcoded variant, but some variants had more than one mutation (Supplemental Figure 1B,C).

### Effects of mutations ot TRO.11 Env on cell entry

To measure the effects of mutations on Env-mediated cell entry, we generated the mutant Env pseudovirus libraries with and without the glycoprotein VSV-G as an additional protein on the virion surface (Figure 2). All virions can infect cells when VSV-G is present, but only virions with functional Env can infect cells when VSV-G is not present. We infected TZM-bl cells, which express Env’s receptor (CD4) and both co-receptors (CCR5 and CXCR4)^31–33^, with each pseudovirus pool. We then calculated cell entry scores for each barcoded variant as the log of the ratio of the change in frequency of that variant relative to the unmutated Env between the conditions with and without VSV-G. As expected, variants with stop codons had highly negative cell entry scores, whereas unmutated variants or ones with only synonymous mutations had cell entry scores of around zero (Supplemental Figure 1D). Variants with amino-acid mutations had functional scores that ranged from zero (“wildtype”-like Env entry) to highly negative (Supplementary Figure 1D). To determine the effects of each individual amino-acid mutation on cell entry, we deconvolved the per-variant cell entry scores using global epistasis models^34,35^.

**Figure 2:**
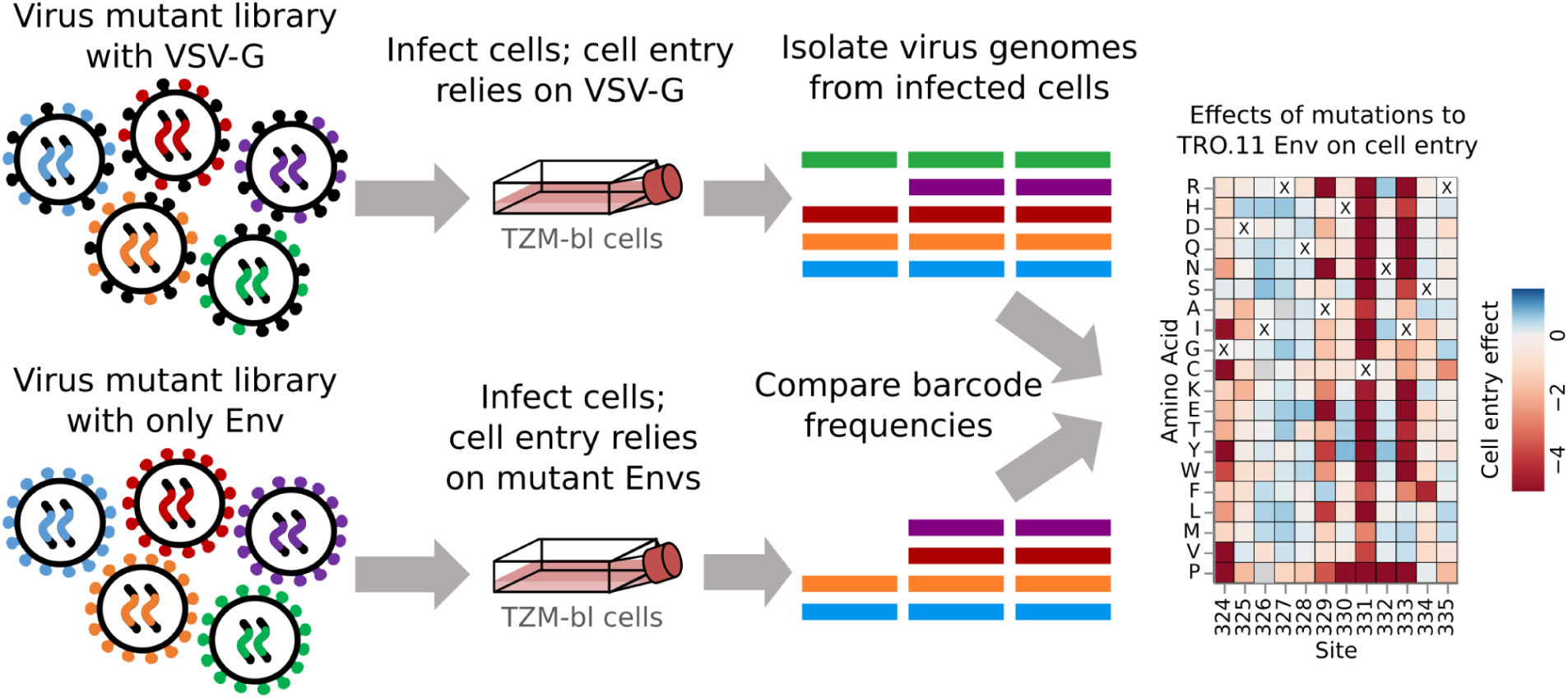
Approach for measuring the effects of mutations on Env-mediated cell entry. Genotype-phenotype linked pseudovirus libraries are produced with and without the addition of the VSV-G glycoprotein. All viruses with VSV-G can enter cells regardless of Env function, but only viruses with a functional Env can enter cells without VSV-G. Genomes are isolated from infected cells in each condition, the barcodes identifying the variants are deep sequenced, and the frequencies of variants between conditions are compared to infer mutation effects on cell entry.

The effects of mutations to the TRO.11 Env on cell entry are shown in Figure 3 (we also suggest the reader examine the interactive version of this heatmap at https://dms-vep.org/HIV_Envelope_TRO11_DMS_3BNC117_10-1074/htmls/TZM-bl_entry_func_effects.html). Sites in the variable loops of the TRO.11 Env (V1 to V5) are tolerant of most mutations. Mutations disrupting N-linked glycosylation motifs (NXS/T where X is any amino acid other than proline) are tolerated for cell entry in most cases, with the notable exceptions of N156 and N262 which have been found to be intolerant of mutations in previous studies (Figure 3)^20,36^. Sites involved in CD4 binding are generally intolerant of mutations, such as most sites at loop D and the CD4 binding loop (Figure 3).

**Figure 3:**
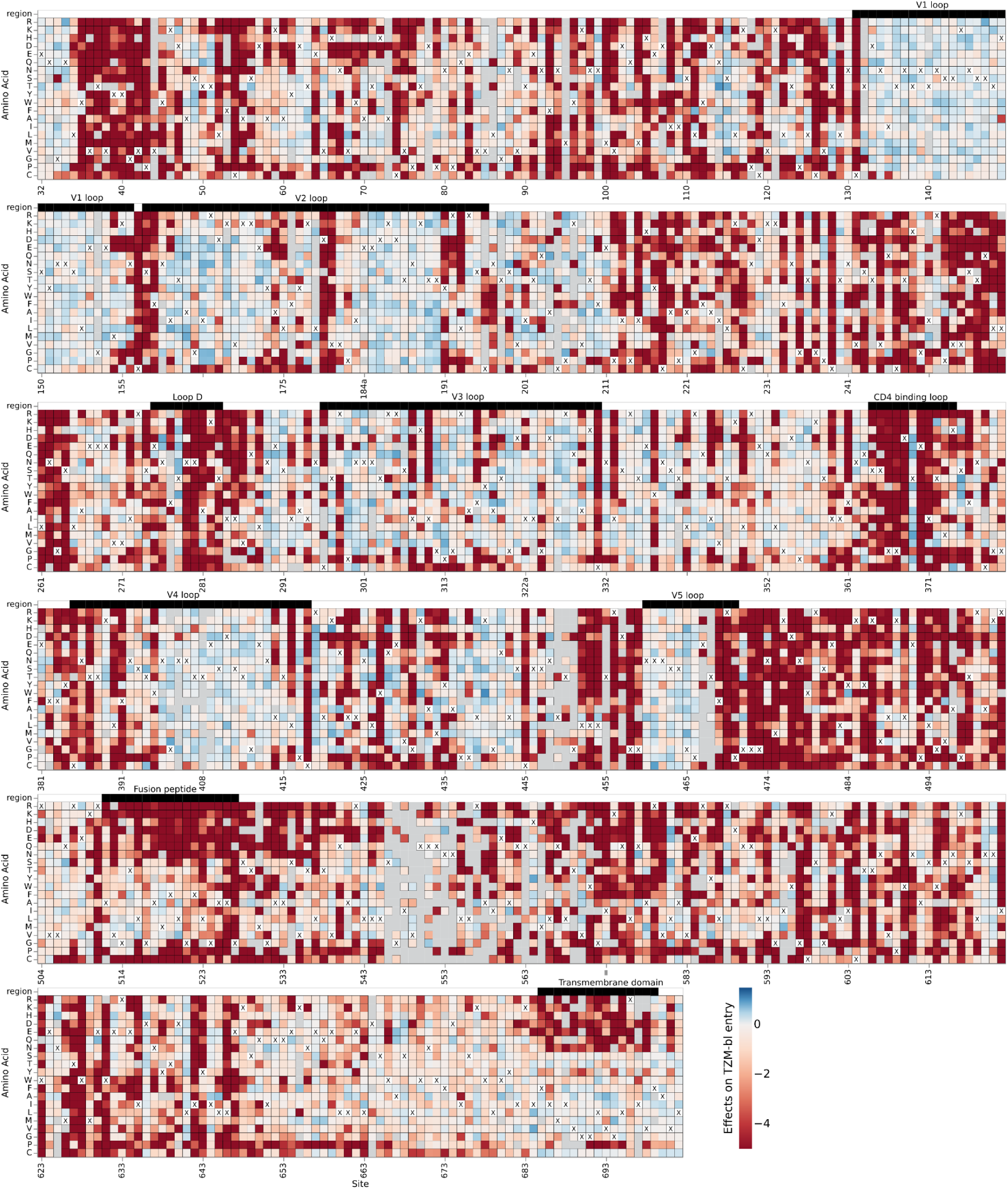
Effects of mutations on TRO.11 Env cell entry. A cell entry effect of zero (white) indicates no effect relative to unmutated TRO.11 Env, negative values (red) indicate impaired entry relative to unmutated TRO.11 Env, and positive values (blue) indicate improved entry relative to unmutated TRO.11 Env. The parental amino-acid identity at each site in TRO.11 Env is labeled with a black “X”. Key regions of Env are denoted with labeled black bars above each row. Mutations with effects that were not well measured in our experiments are colored gray. See https://dms-vep.org/HIV_Envelope_TRO11_DMS_3BNC117_10-1074/htmls/TZM-bl_entry_func_effects.html for an interactive version of this heatmap.

### Measuring effects of Env mutations on escape from antibody neutralization

To measure the effects of mutations on antibody-neutralization, we mixed the Env pseudovirus libraries with a small amount of VSV-G pseudotyped viruses carrying known barcodes, which act as non-neutralized standards that enable conversion of sequencing counts to the absolute fraction of each variant that is neutralized (Figure 4)^18^. The Env mutant library with the VSV-G standard was then incubated with a range of antibody concentrations before being infected into TZM-bl cells, and barcodes were extracted and deep sequenced (Figure 4). The extent of neutralization of each Env variant at each antibody concentration was computed by comparing the barcode counts for that Env to those for the VSV-G standard in the conditions with and without antibody (Figure 4). The overall effects of each mutation on antibody neutralization were then computed from these values using a previously described biophysical model^37^.

**Figure 4:**
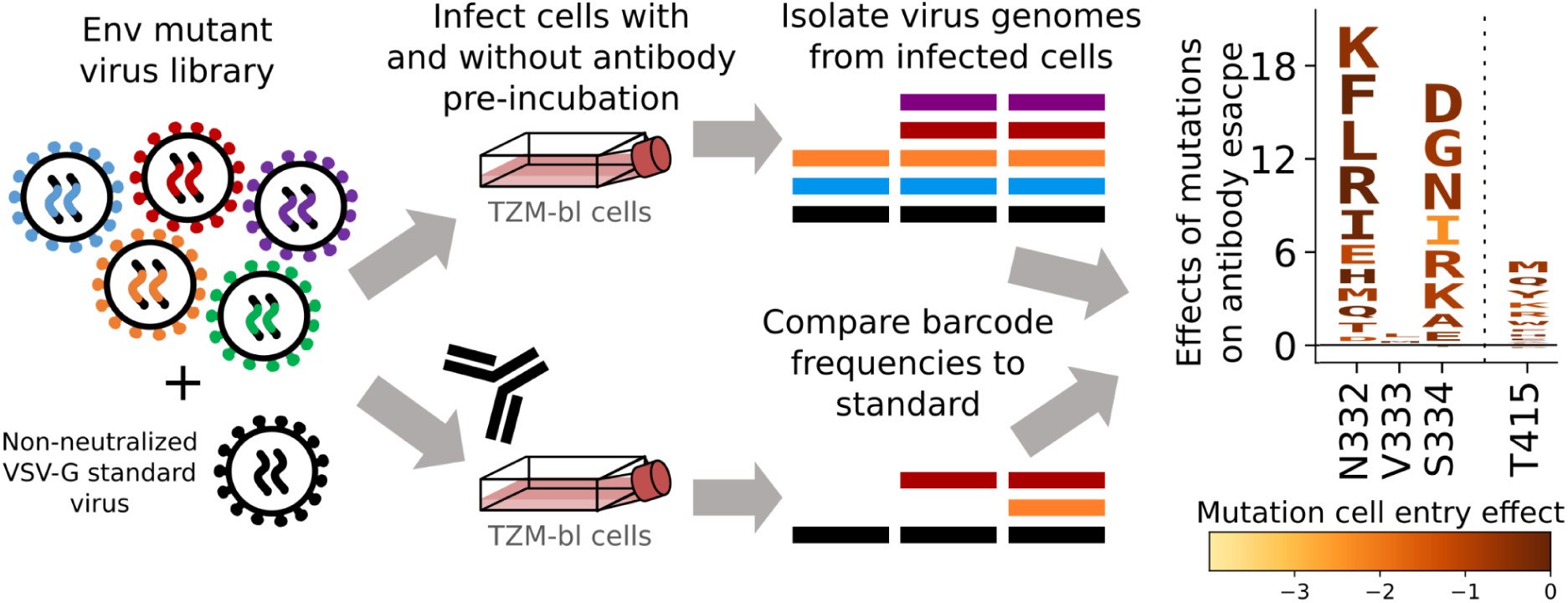
Measuring effects of Env mutations on antibody neutralization. Genotype-phenotype linked Env mutant libraries are mixed with a small amount of separately produced VSV-G pseudotyped standard viruses with known barcodes (VSV-G is not neutralized by anti-Env antibodies). The library plus standard is then incubated with various concentrations of antibody (or no antibody for the control condition) for one hour prior to infecting cells. Genomes are isolated from infected cells in each condition, barcodes corresponding to each variant and the standards are deep sequenced, and the frequency change of each variant between the no-antibody control and the antibody conditions is analyzed to determine the effect of each mutation on neutralization. The data are represented in logo plots, with the height of each letter proportional to how much it escapes antibody neutralization. Letters are colored by the effect of that mutation on Env-mediated cell entry.

### Escape from antibody 10-1074

We used the workflow described above to measure how mutations affected neutralization by antibody 10-1074^3^ for both the TRO.11 Env libraries described above and our previously described BF520 libraries^26^. Plots showing the effects on neutralization by all tolerated mutations to each Env are shown in Supplementary Figures 3 and 4 and in interactive form at https://dms-vep.org/HIV_Envelope_TRO11_DMS_3BNC117_10-1074/htmls/10-1074_mut_effect.html and https://dms-vep.org/HIV_Envelope_BF520_DMS_3BNC117_10-1074/htmls/10-1074_mut_effect.html. Below we focus our analysis on just the sites where mutations have an appreciable effect on 10-1074 neutralization for at least one of the Envs.

Mutations that disrupt the N332 glycosylation motif cause a substantial amount of escape from neutralization by antibody 10-1074 for both BF520 Env and TRO.11 Env, but the effects of mutations at nearby sites slightly differ between the two Envs (Figure 5). Mutations in and near the co-receptor binding site spanning residues 324 to 327^11^ cause escape in both strains, but the effects of specific mutations at these sites differ between strains (Figure 5A,B). Mutations away from negatively charged amino acids D or E at site 325 caused escape for both strains (Figure 5A,B). Mutations at site R327 were measured to cause escape for TRO.11 Env, but not BF520 Env. This may be due to TRO.11 Env being tolerant of most mutations at this site (Figure 3), while BF520 Env is tolerant of almost no mutations at this site (Supplemental Figure 2).

**Figure 5:**
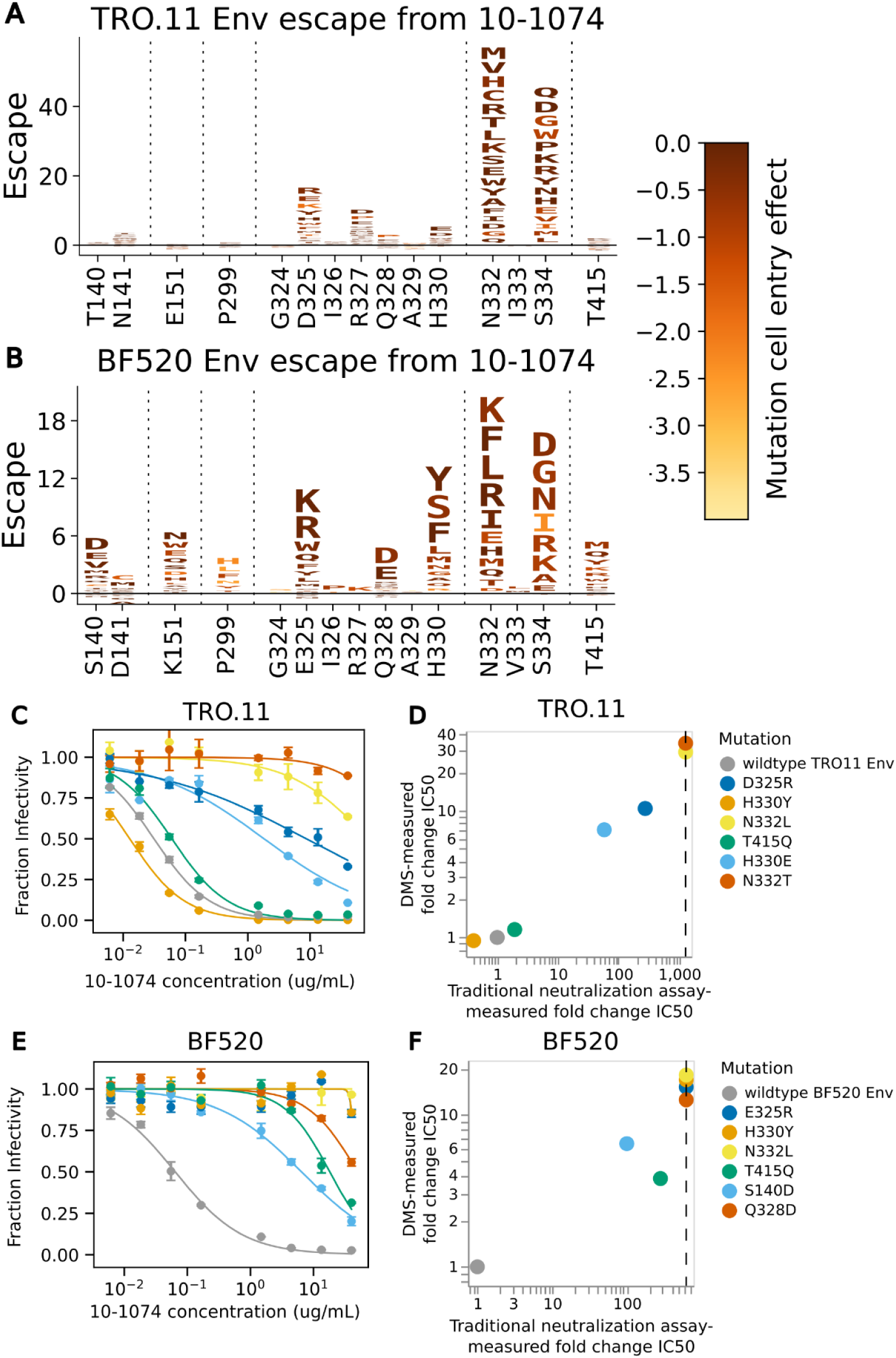
Escape from antibody 10-1074. (A) Logoplot showing effects of TRO.11 Env mutations on escape from antibody 10-1074. The height of each letter represents the effect of that amino-acid mutation on antibody neutralization, with positive heights (letters above the zero line) indicating mutations that cause escape, and negative heights (letters below the zero line) indicating mutations that increase neutralization. Letters are colored by the effect of that mutation on pseudovirus cell entry, with yellow corresponding to impaired cell entry and brown corresponding to neutral effects on cell entry. Only key sites are shown. See https://dms-vep.org/HIV_Envelope_TRO11_DMS_3BNC117_10-1074/htmls/10-1074_mut_effect.html for an interactive version of the escape map. (B) Logoplot showing effects of BF520 Env mutations on escape from antibody 10-1074. See https://dms-vep.org/HIV_Envelope_BF520_DMS_3BNC117_10-1074/htmls/10-1074_mut_effect.html for an interactive version of the escape map. (C) Neutralization curves for unmutated TRO.11 Env pseudoviruses and selected single mutants. (D) Scatter plot of TRO.11 Env fold-change IC50s measured by deep mutational scanning versus measured in the neutralization assays. Vertical dotted lines represent the limit of detection of the neutralization assays. See methods section “Analysis of the effects of mutations on HIV Env escape from neutralization by antibodies” for details on how deep mutational scanning measured escape values are converted to deep mutational scanning measured IC50 values (E) Neutralization curves for unmutated BF520 Env pseudoviruses and selected single mutants. (F) Scatter plot of BF520 Env mutant fold-change IC50s measured by deep mutational scanning versus measured in the neutralization assays.

Mutations to negatively charged amino acids D or E were measured to cause escape at site Q328 for BF520 Env, but have a small relative effect on escape for TRO.11 Env (Figure 5A,B). Any mutations at H330 were measured to cause high escape for BF520 Env, but only mutations to negatively charged amino acids (D or E) were measured to cause high escape relative to other mutations for TRO.11 Env (Figure 5A, B). There were additional sites with escape mutations for BF520 Env, such as sites 140, 141, 151, 299, and 415 (Figure 5A,B).

Traditional neutralization assays were performed with pseudoviruses with single mutations to BF520 Env or TRO.11 Env to validate the deep mutational scanning escape maps (Figure 5C-F). Deep mutational scanning measured escape was well-correlated with traditional neutralization assay-measured fold change IC50s (Figure 5D,F). These validation assays also confirmed the Env-specific escape shown by the deep mutational scanning: for instance, H330Y and T415Q caused escape in BF520 but not TRO.11 (Figure 5C-F).

### Escape from antibody 3BNC117

We used the TRO.11 Env libraries and the workflow described above to measure how mutations affect neutralization by antibody 3BNC117^4^. We also re-analyzed the effects of BF520 Env mutations on neutralization by antibody 3BNC117 from our prior deep mutational scanning study^26^. Plots showing the effects on neutralization by all tolerated mutations to each Env are shown in Supplementary Figures 5 and 6 and in interactive form at https://dms-vep.org/HIV_Envelope_TRO11_DMS_3BNC117_10-1074/htmls/3BNC117_mut_effect.html and https://dms-vep.org/HIV_Envelope_BF520_DMS_3BNC117_10-1074/htmls/3BNC117_mut_effect.html. Below we focus our analysis on just the sites where mutations have an appreciable effect on 3BNC117 neutralization for at least one of the Envs.

Mutations measured to cause escape from neutralization by antibody 3BNC117 differ greatly between BF520 Env and TRO.11 Env (Figure 6). Some sites where mutations have different effects on neutralization by 3BNC117 between BF520 Env and TRO.11 Env differ in their tolerance for mutations between the two Envs. At loop D, TRO.11 Env is intolerant of any mutations at sites 279 and 280, and intolerant of most mutations at site 281 (Figure 3). Mutations are tolerated at these sites in BF520 Env (Supplemental Figure 2) and mutations at these sites were measured to cause escape from antibody 3BNC117 for BF520 Env (Figure 6B), but were not measured to cause escape for TRO.11 Env because those mutants were no longer able to enter cells. Conversely, mutations to negatively charged amino acids at sites 304, 308, and 318 in the V3 loop cause escape from 3BNC117 for TRO.11 Env but not BF520 Env (Figure 6A,B); TRO.11 Env is tolerant of negatively charged amino acids in this region (Figure 3), but BF520 Env is not (Supplemental Figure 2). Some mutations at site 456 were measured to cause escape from BF520 Env (Figure 6B), but TRO.11 Env is intolerant of any mutations at site 456 (Figure 2).

**Figure 6:**
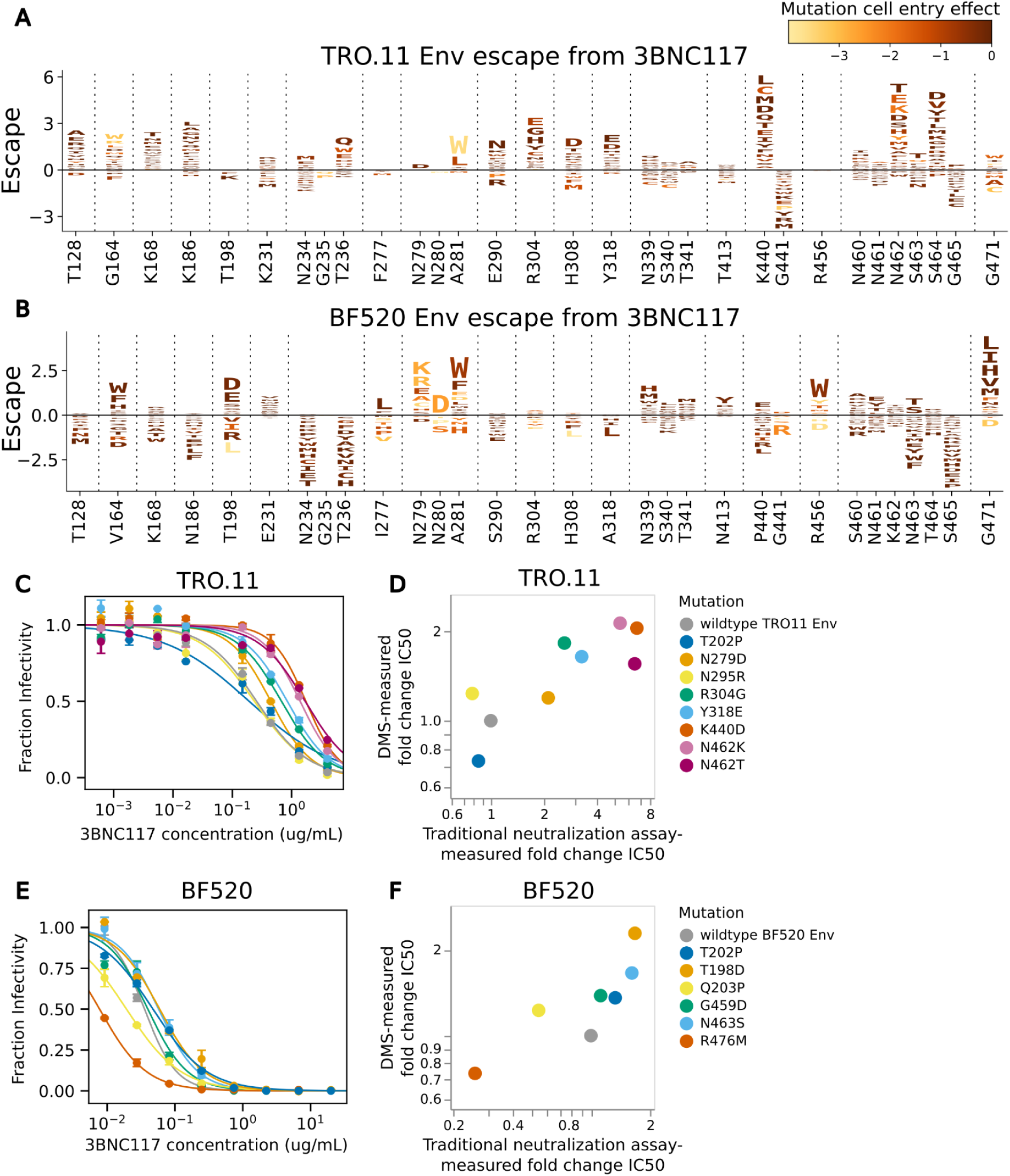
Escape from 3BNC117. (A) Logoplot showing effects of TRO.11 Env mutations on escape from antibody 3BNC117. The height of each letter represents the effect of that amino-acid mutation on antibody neutralization, with positive heights (letters above the zero line) indicating mutations that cause escape, and negative heights (letters below the zero line) indicating mutations that increase neutralization. Letters are colored by the effect of that mutation on pseudovirus cell entry, with yellow corresponding to impaired cell entry and brown corresponding to neutral effects on cell entry. Only key sites are shown. See https://dms-vep.org/HIV_Envelope_TRO11_DMS_3BNC117_10-1074/htmls/3BNC117_mut_effect.html for an interactive version of the escape map. (B) Logoplot showing effects of BF520 Env mutations on escape from antibody 3BNC117. See https://dms-vep.org/HIV_Envelope_BF520_DMS_3BNC117_10-1074/htmls/3BNC117_mut_effect.html for an interactive version of the escape map. (C) Neutralization curves for unmutated TRO.11 Env pseudoviruses and selected single mutants. (D) Scatter plot of TRO.11 Env fold-change IC50s measured by deep mutational scanning versus measured in the neutralization assays. See methods section “Analysis of the effects of mutations on HIV Env escape from neutralization by antibodies” for details on how deep mutational scanning measured escape values are converted to deep mutational scanning measured IC50 values. (E) Neutralization curves for unmutated BF520 Env pseudoviruses and selected single mutants. (F) Scatter plot of BF520 Env mutant fold-change IC50s measured by deep mutational scanning versus measured in the neutralization assays. The BF520 Env mutant neutralization curves and the BF520 Env deep mutational scanning measured effects of mutations on escape from antibody 3BNC117 shown in this figure are previously published^26^.

However, many sites where the effects of mutations on 3BNC117 neutralization differ between the Envs have similar tolerance of mutations. Mutations at site 164 cause some escape for both Envs, but mutations at nearby sites 128, 168, and 186 cause escape for TRO.11 Env, but not BF520 Env despite similar levels of mutational tolerance in this region (Figure 6A,B, Supplemental Figure 2,3). Any mutations that disrupt the N234 glycosylation motif cause better neutralization by 3BNC117 for BF520 Env, but specific mutations at N234 and T236 cause escape from 3BNC117 for TRO.11 Env (Figure 6A,B). Any mutations at site 440 cause escape for TRO.11 Env, but not for BF520 Env despite being tolerated (Figure 6A,B, Supplemental Figure 2,3). At the V5 loop, TRO.11 Env has two tandem N-linked glycosylation motifs, at N461 and N462 (Figure 6A). Disruption of the N462 glycosylation motif by any mutations causes escape for TRO.11 Env, possibly by making a glycosylation of N461 more likely (Figure 6A). BF520 Env has a single N-linked glycosylation motif at the V5 loop at N463 (Figure 6B). Disruption of this glycosylation motif causes better neutralization by 3BNC117 for BF520 Env, but mutations N463T and N463S that shift the glycosylation motif to N461 cause escape from 3BNC117 for BF520 Env (Figure 6B). Mutations at site 471 cause a high amount of escape for BF520 Env, but not TRO.11 Env, despite TRO.11 Env being tolerant of some mutations at this site (Figures 2, 6A,B).

We performed traditional neutralization assays to validate the deep mutational scanning 3BNC117 escape maps for TRO11 Env, and re-analyzed neutralization assays using 3BNC117 from our previous study^26^ for BF520 Env. The deep mutational scanning measured escape was well-correlated with traditional neutralization assay-measured fold change IC50s (Figure 6D,F).

### Differences in the effects of mutations on neutralization are not explained by parental amino acid conservation between Env strains

One hypothesis is that the sites where mutations have different effects on antibody escape between the Envs are those sites where the parental amino-acid identities differ between the Envs. However, closer investigation reveals that conservation of the parental amino acid between BF520 Env and TRO.11 Env is not a strong determinant of whether the effects of mutations on neutralization at that site are consistent between Envs (Figure 7).

**Figure 7:**
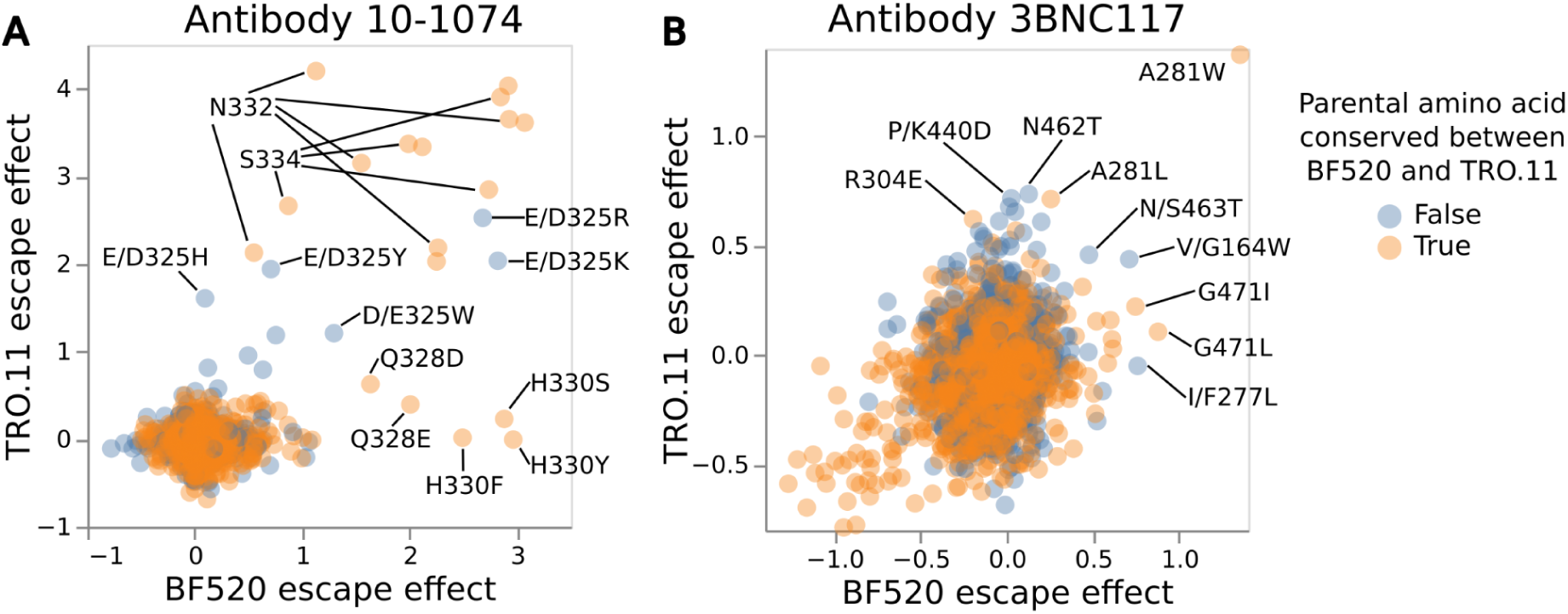
Effects of mutations on neutralization of the TRO.11 versus BF520 Env annotated by whether the mutated site is conserved between the Envs. Scatter plots of the deep mutational scanning measured effects of each mutation on escape from 10-1074 (A) or 3BNC117 (B) for the BF520 Env (x-axis) versus the TRO.11 Env escape (y-axis). Mutations are colored by whether the parental amino acid identity is conserved between BF520 Env and TRO.11 Env. Key mutations are labeled; when the parental amino acid identity is not conserved between BF520 Env and TRO.11 Env then the labels give the BF520 Env identity as the first wildtype amino acid and the TRO.11 Env identity as the second wildtype amino acid. See https://dms-vep.org/HIV_Envelope_TRO11_DMS_3BNC117_10-1074/notebooks/strain_escape_comparison.html for interactive versions of these plots. BF520 Env deep mutational scanning measured effects of mutations on escape from antibody 3BNC117 shown in this figure are previously published^26^.

Consistency between Envs for the effects of mutations on escape from 10-1074 cannot be predicted by parental amino acid conservation (Figure 7A). Any mutations at conserved sites N332 or S334 of the N332 glycosylation motif cause escape from 10-1074 for both Envs (Figure 7A). However, there are mutations at conserved sites Q328 and H330 that only cause escape from 10-1074 for BF520 Env and not TRO.11 Env (Figure 7A). Mutations at non-conserved site D/E325 cause escape for both Envs (Figure 7A), but it is worth noting D and E are both negatively charged amino acids and mutations there likely have similar biophysical effects for both Envs.

Consistency between Envs for the effects of mutations on escape from 3BNC117 also cannot be predicted by parental amino acid conservation (Figure 7B). Mutations at conserved site A281 were measured to cause escape from 3BNC117 for both Envs (Figure 7B), but mutations at conserved site G471 only caused appreciable escape from 3BNC117 for BF520 Env and not TRO.11 Env (Figure 7B). Mutations at non-conserved site K/P440 cause escape from 3BNC117 for TRO.11 Env but not for BF520 Env (Figure 7B), while mutations at nonconserved site G/V164 and mutations S/N463T cause escape for both Envs from 3BNC117 (Figure 7B).

## Discussion

We used a lentiviral vector deep mutational scanning platform to measure the effects of mutations on TRO.11 Env cell entry and escape from neutralization by antibodies 10-1074 and 3BNC117. We previously used this platform to measure the effects of mutations on BF520 Env cell entry and escape from neutralization by 3BNC117, and in this study we also measured the effects of mutations on BF520 Env escape from 10-1074. Together, these datasets provide detailed maps of the effects of mutations on cell entry and escape from two clinically relevant broadly neutralizing antibodies for two HIV Envs from different clades of HIV.

Overall, we found that the escape mutations from antibody 10-1074 were largely similar (although still not identical) between the two Envs, but the escape mutations for 3BNC117 were very different between the Envs. Neutralization by 10-1074 is highly dependent on the N332 glycan and a short stretch of nearby amino acids^3,38^, and we found mutations at these sites cause the most escape from 10-1074 relative to other sites in both Envs. On the other hand, neutralization by 3BNC117 is dependent on antibody-protein interactions spread across Env^12–14^, and we found that sites where mutations cause the most escape from 3BNC117 are not consistent between Envs. These differences can be partially explained by general differences in tolerance of mutations in Env regions between the Envs and glycosylation patterns (see below), but for some sites the reason for the difference in escape effects between the Envs is unclear.

Some of the differences between Envs for the effects of mutations on escape from neutralization by 10-1074 or 3BNC117 can be explained by differences in tolerance of mutations between the Envs. In particular, the TRO.11 Env is generally more tolerant of mutations to negatively charged amino acids in the V3 loop, and this results in differences in escape between Envs for both antibodies. Mutations H330D/E cause escape from 10-1074 for TRO.11 Env but are not tolerated for BF520 Env, and mutations to D and E at sites 304, 308, and 318 cause escape from 3BNC117 for TRO.11 Env but are not tolerated for BF520 Env. However, this observation is not entirely straightforward; mutations Q328D/E are tolerated by both Envs, but for some reason only cause escape from 10-1074 for BF520 Env. Additional sites with parental amino acids conserved between the Envs have differences in effects on escape as well. These examples highlight the difficulty in predicting HIV Env mutations that escape antibodies: strains differ greatly in tolerance of mutations across sites, and the effects of mutations on escape can still differ between strains at conserved sites with similar tolerance of mutations. We speculate that 3BNC117’s interaction with residues across several regions of Env allow more chances for systemic differences in Env tolerance of mutations or glycosylations to affect escape, leading to more striking differences in escape between Envs than for 10-1074.

Our results underscore the importance of the presence or absence of glycosylations at certain sites for HIV Env neutralization by antibodies. Antibody 10-1074’s reliance on the presence of the N332 glycosylation for neutralization is well-documented^3,14,38^ and demonstrated again in our study. More subtly, we found that the presence of a glycosylation at N460 or N461, but not at N462, causes escape from antibody 3BNC117 for both the TRO.11 and BF520 Envs. However, the effect of mutations at some other glycosylation motifs on escape differs among Envs due to mutation tolerance and glycosylation pattern differences. Prior deep mutational scanning of BG505 Env showed escape from 3BNC117 was caused by mutations N197S/T by shifting a glycosylation motif from N197 to N195^14^, but N197S/T are not tolerated by BF520 Env and N197S/T does not introduce a new glycosylation motif for TRO.11 Env because it has S195 rather than N195. The prior deep mutational scanning of BG505 Env also showed no escape from 3BNC117 for mutations introducing a glycosylation motif at site 461, although BG505 Env would retain a glycosylation motif at N462 without additional mutations^14^.

The two strains we use here, BF520 Env and TRO.11 Env, represent just a small subset of the diversity of HIV. Nonetheless, our results are sufficient to begin to shed some light on the question of how consistent (and hence predictable) antibody-escape mutations are across Envs from different viral strains. For one of the two antibodies we studied, escape mutations were quite consistent even across two Envs from different strains, but for the other antibody they were quite diverged. This fact suggests that the ability to prospectively identify escape mutations in a way that can inform predictive models applicable to many viral strains may vary across antibodies. Future studies using mutant libraries spanning more clades of HIV and antibodies could shed further light into how antibody-escape mutations vary across viral strains.

## Acknowledgments

We thank Dr. Michel Nussenzweig and Marina Caskey for a gift of 3BNC117 and 10-1074 IgGs. This work was supported by the NIAID/NIH under grants R01AI140891 and U01AI169385. JDB is an Investigator of the Howard Hughes Medical Institute. This research also was supported by the Genomics & Bioinformatics Shared Resource (RRID:SCR_022606), Flow Cytometry Shared Resource (RRID:SCR_022613), and Scientific Computing (NIH grants S10-OD-020069 and S10-OD-028685) of the Fred Hutch/University of Washington/Seattle Children’s Cancer Consortium (P30 CA015704).

## Methods

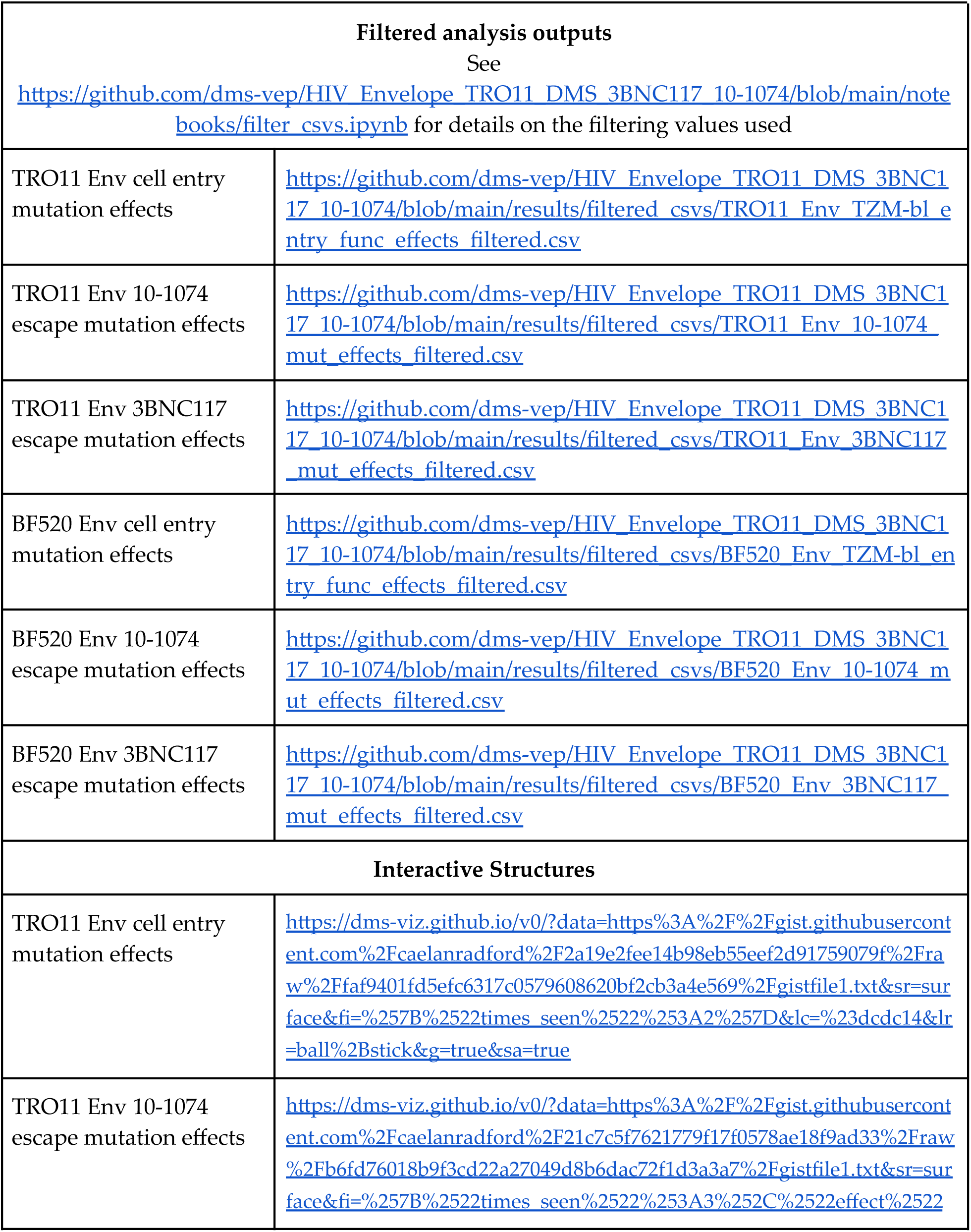
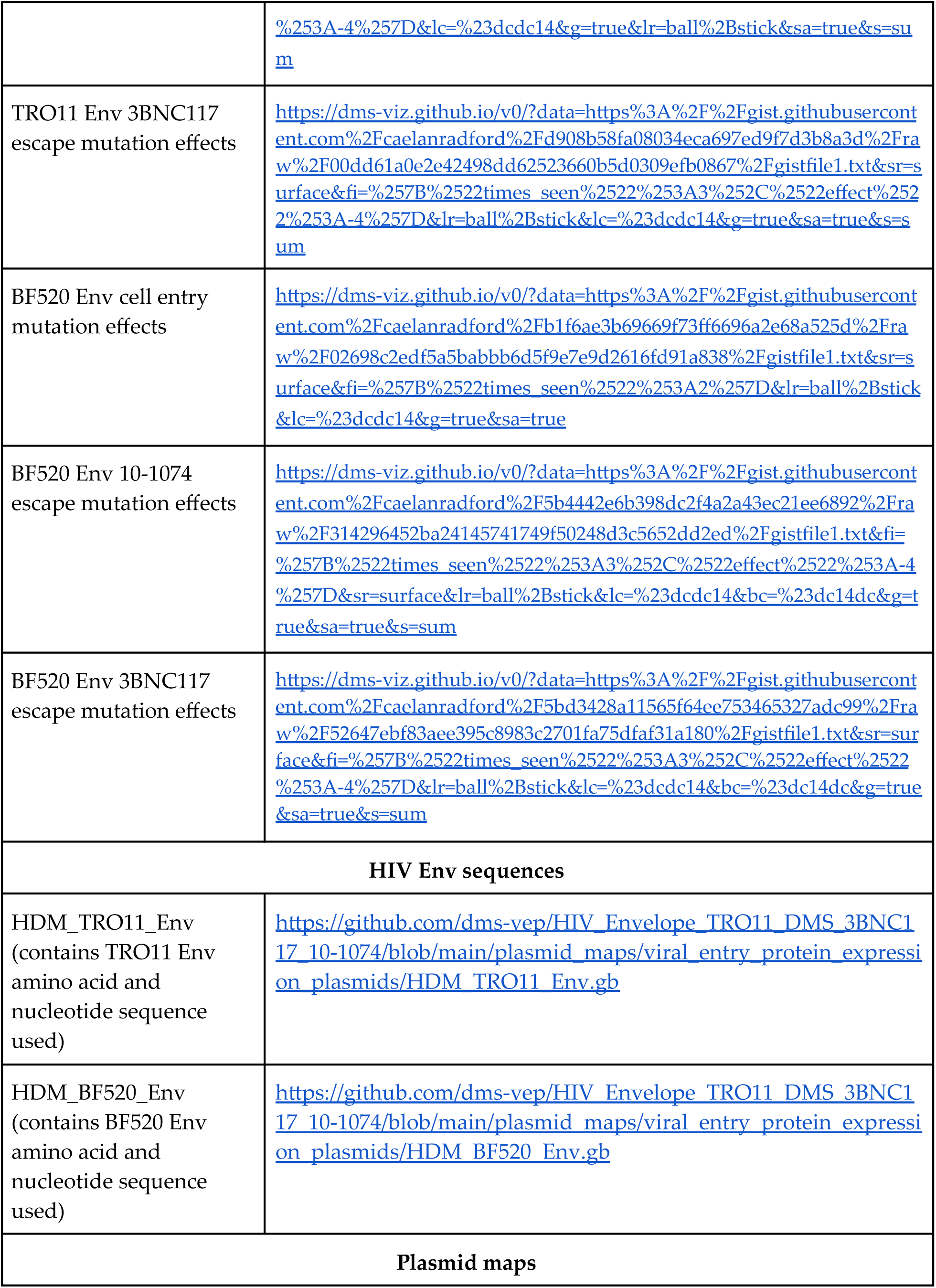
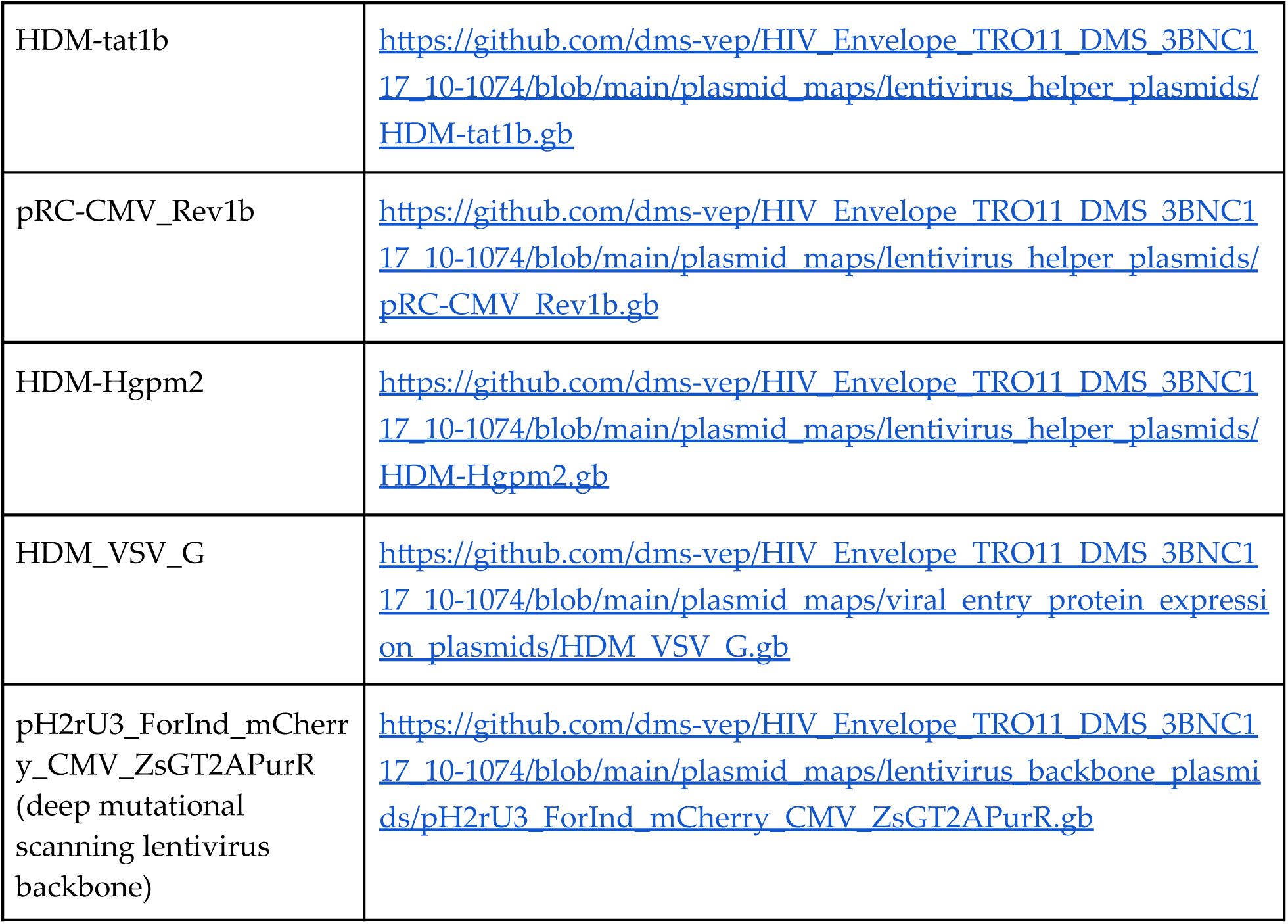
Table of key outputs and resources.

### Lentivirus deep mutational scanning platform

We used a lentivirus-based deep mutational scanning platform that we have previously described^18,26^. The platform generates tens of thousands of mutant lentivirus pseudovirus variants with a genotype-phenotype link between the mutant viral entry protein on the surface of the virions and a random nucleotide barcode in the lentivirus genome. The process for producing the genotype-phenotype linked pseudovirus libraries is described in detail previously^18,26^. See https://github.com/dms-vep/HIV_Envelope_TRO11_DMS_3BNC117_10-1074/blob/main/plasmid_maps/lentivirus_backbone_plasmids/pH2rU3_ForInd_mCherry_CMV_ZsGT2APurR.gb for a plasmid map containing the lentivirus genome with mCherry in the position into which viral entry protein mutants are cloned.

### Design of and cloning of HIV Env mutant libraries

The BF520 Env mutant libraries were cloned and described in a previous study^26^. See Radford et al.^26^ for details. The BF520 Env mutant libraries were cloned using a codon tiling primer mutagenesis strategy to target well-tolerated mutations and a high mutation rate per Env mutant (∼3 mutations per BF520 Env mutant). The TRO.11 Env mutant libraries were generated for this study. Since we did not have prior deep mutational scanning data for the effects of mutations on TRO.11 Env cell entry, the TRO.11 Env mutant libraries are site saturation mutagenesis libraries targeting all mutations in the Env ectodomain with a lower mutation rate per Env mutant (∼1 mutation per TRO.11 Env mutant).

We use the HXB2 numbering scheme for sites in Env throughput this manuscript^44^. For the TRO.11 Env mutant libraries, we aimed to generate all amino acid mutations at all sites in the TRO.11 Env ectodomain from sites 31 to 702 during mutagenesis. We also aimed to generate stop codons at alternating sites from sites 31 to 192, to be used as negative controls as non-functional mutants. We ordered single site saturation mutagenesis libraries from Twist Bioscience. The Twist quality control report indicated there were 22 sites that failed mutagenesis and were not included in the delivered library.

We sought to separately mutagenize the failed TRO.11 Env sites through NNS primer mutagenesis. To do this, we ordered forward and reverse primers with an NNS sequence at the failed codon in the middle of the primer and 20 base pair overhangs on each side. For each site, we did separate forward and reverse mutagenesis reactions. We used the following conditions for the forward reactions: PCR mix: 21 µL H2O, 1.5 µL 10 µM forward mutagenesis primer for the site, 1.5 µL 10 µM reverse primer (Read1_rev, AGAGCGTCGTGTAGGGAAAGAGTG), 1 µL 10ng/µL TRO.11 Env template (HDM_TRO11_Env, see https://github.com/dms-vep/HIV_Envelope_TRO11_DMS_3BNC117_10-1074/blob/main/plasmid_maps/viral_entry_protein_expression_plasmids/HDM_TRO11_Env.gb), 25 µL 2x KOD Hot Start Master Mix (Sigma-Aldrich, Cat. No. 71842). Cycling conditions: (1) 95C/2min (2) 95C/20sec (3) 55 C/10sec (4) 70C/60 sec (5) Return to Step 2 x24. We used the following conditions for the reverse reactions: PCR mix: 21 µL H2O, 1.5 µL 10 µM reverse mutagenesis primer for the site, 1.5 µL 10 µM foward primer (VEP_amp_for, CAGCCGAGCCACATCGCTC), 1 µL 10ng/µL TRO.11 Env template (HDM_TRO11_Env), 25 µL 2x KOD Hot Start Master Mix. Cycling conditions for the reverse reactions were the same as for the forward reactions. The products were DpnI treated overnight, and then joined using a joining PCR with the following conditions: PCR mix: 19 µL H2O, 1.5 µL forward mutagenized fragment, 1.5 µL reverse mutagenized fragment, 1.5 µL 10 µM foward primer (VEP_amp_for), 1.5 µL 10 µM reverse primer (Read1_rev), 25 µL 2x KOD Hot Start Master Mix. Cycling conditions: (1) 95C/2min (2) 95C/20sec (3) 58 C/10sec (4) 70C/60 sec (5) Return to Step 2 x14. The joined PCR products for each site were Ampure XP bead (Beckman Coulter, Cat. No. A63881) cleaned, quantified using a Qubit Fluorometer (ThermoFisher), and pooled at an even ratio. The pooled NNS mutagenized fragments were then barcoded using the barcoding PCR described below.

We linearized and amplified the Twist mutagenized fragments using a PCR with the following conditions: PCR mix: 17 µL H2O, 5 µL of Twist mutagenized fragment pool, 1.5 µL forward primer (VEP_amp_for), 1.5 µL reverse primer (Read1_rev), 25 µL KOD Hot Start Master Mix. Cycling conditions: (1) 95C/2min (2) 95C/20sec (3) 58 C/10sec (4) 70C/60 sec (5) Return to Step 2 x14. We gel extracted the product and Ampure bead cleaned the product.

We then separately barcoded the linearized mutagenized fragments from the Twist pool and from the NNS mutagenesis pool using the following conditions for each: 30 ng of linearized mutagenized fragments, 1.5 µL 5 µM forward primer (VEP_amp_for), 1.5 µL 5 µM ForInd_AddBC_XbaI (barcoding primer: catttctctctcgaaTCTAGANNNNNNNNNNNNNNNNAGATCGGAAGAGCGTCGTGTAGGGAAAG), 15 µL KOD Hot Start Master Mix, filled to 30 µL total with H2O. Cycling conditions: Cycling conditions: (1) 95C/2min (2) 95C/20sec (3) 70C/1sec (4) 50C/10sec, cooling at 0.5/sec (5) 70C/120sec (6) Return to Step 2 x9. The barcoded products were gel extracted, Ampure bead cleaned (eluting into H2O), and quantified using Qubit. The barcoded Twist mutagenized fragments and the barcoded NNS mutagenized fragments were pooled by mass at the ratio of successful Twist mutagenized sites to NNS mutagenized sites.

We digested the recipient lentivirus genome (pH2rU3_ForInd_mCherry_CMV_ZsGT2APurR.gb, see link in section above) with MluI and XbaI, gel extracted the digested vector, Ampure bead cleaned (eluting into H2O), and quantified using Qubit. We used a 2:1 insert to vector ratio in a 1 hour HiFi assembly reaction using NEBuilder HiFi DNA Assembly kit (NEB, Cat. No. E2621), Ampure bead cleaned the product (eluting into 20 µL H2O), and performed ten separate electroporations per replicate TRO.11 Env library by using 2 µL of the cleaned HiFi product to transform 20 µL of 10-beta electrocompetent E. coli cells (NEB, C3020K). This yielded >5 million CFUs per replicate library. We plated each electroporation on a 15 cm LB+ampcicillin plate, incubated the plates overnight at 37C, and scraped the plates the next day. The pooled bacteria for each replicate library were diluted to 15 OD600 and then used in five separate 5 mL minipreps (QIAprep Spin Miniprep Kit, Cat. No. 27106X4) each. The leftover bacteria for each replicate library were spun down in pellets and stored at −20C.

### Generation of cell line-stored HIV TRO.11 Env mutant libraries

Generation of genotype-phenotype linked pseudoviruses requires a multi-step process that includes 1) production of a diverse pool of VSV-G pseudotyped pseudoviruses carrying the barcoded Env mutants, 2) infection of 293T-rtTA cells at a low multiplicity of infection, such that each infected cell is infected with only one lentivirus genome, and 3) enrichment of infected 293T-rtTA cells using puromycin selection. This process was completed and described for the BF520 Env mutant libraries in a prior study^26^. See Radford et al.^26^ for details. The following section described the conditions used for this process for the TRO.11 Env mutant libraries, which were nearly identical.

To not bottleneck the mutant barcoded variants during the production of the pool of VSV-G pseudotyped viruses, we aimed to maintain a high diversity at this step by producing many more viruses than the intended eventual mutant library size of ∼80,000 barcoded variants. The day before transfection, we plated 800,000 293T cells per well in six well plates. For each replicate TRO.11 Env mutant library, we transfected six separate six well plates. We used BioT (Bioland Scientific) as the transfection reagent, following the manufacturer’s recommendations. We transfected each well with 1 microgram of the lentivirus genome with the barcoded TRO.11 Env mutants and 250 nanograms each of a HIV Tat expressing plasmid (HDM-tat1b, see https://github.com/dms-vep/HIV_Envelope_TRO11_DMS_3BNC117_10-1074/blob/main/plasmid_maps/lentivirus_helper_plasmids/HDM-tat1b.gb), a HIV Rev expressing plasmid (pRC-CMV_Rev1b, see https://github.com/dms-vep/HIV_Envelope_TRO11_DMS_3BNC117_10-1074/blob/main/plasmid_maps/lentivirus_helper_plasmids/pRC-CMV_Rev1b.gb), a HIV Gag-Pol expressing plasmid (HDM-Hgpm2, see https://github.com/dms-vep/HIV_Envelope_TRO11_DMS_3BNC117_10-1074/blob/main/plasmid_maps/lentivirus_helper_plasmids/HDM-Hgpm2.gb), and a VSV-G expressing plasmid (HDM_VSV_G, see https://github.com/dms-vep/HIV_Envelope_TRO11_DMS_3BNC117_10-1074/blob/main/plasmid_maps/viral_entry_protein_expression_plasmids/HDM_VSV_G.gb). 48 hours after transfection, the supernatants for each replicate library were pooled, aliquoted in 1 mL aliquots, and stored at −80C. To obtain an infectious units per mL estimate for each virus, we titrated these viruses using flow cytometry to estimate ZsGreen percent positivity of cells infected by dilutions of each virus^45^.

We aimed to make TRO.11 Env barcoded variant libraries with around 80,000 variants per replicate library. To do this, we followed the previously described method for the BF520 Env libraries^26^ exactly, but targeted 80,000 variants rather than the 40,000 variants for the BF520 Env libraries. Briefly, we infected 293T-rtTA cells in six well plates, aiming to infect at a 0.005 multiplicity of infection. At the time of infection, we also counted cells in several wells. Two days later, we used flow cytometry and the cell count per well at the time of infection to determine the actual multiplicity of infection obtained, and then pooled wells to obtain 80,000 infected cells for each replicate library based on the number of initially infected cells in each well.

24 hours later, we added 0.75 micrograms per mL of puromycin to the pooled cells to select for transduced cells, which should be resistant to puromycin. The cells for each replicate library were expanded under puromycin selection and eventually frozen in 1 mL aliquots of 20 million cells in tetracycline-negative heat-inactivated fetal bovine serum (Gemini Bio, Cat. No. 100-800) with 10% DMSO. These aliquots of cells were then ready to be thawed and expanded to produce genotype-phenotype linked viruses as described in the following sections.

### Production of genotype - phenotype linked HIV Env mutant virus libraries

Genotype-phenotype linked pseudoviruses can be produced from the cell line-stored Env libraries by transfecting each library cell line with the lentiviral helper plasmids as described previously^18,26^. To produce these viruses, we plated 150 million cells per flask in five layer flasks, which we transfected 24 hours later using BioT, using 50 ug of each lentivirus helper plasmid (Gag-Pol, Rev, and Tat) mixed with 225 µL of BioT mixed and 7.5 mL of DMEM. We added 100 ng/mL of doxycycline to induce Env expression at the time of transfection. 48 hours later, we filtered the supernatant through a .45 µM SFCA filter (Nalgene, Cat. No. 09-740-44B). We concentrated the filtered virus using ultracentrifugation with a 20% sucrose cushion at 100,000 g for one hour. We resuspended the viruses in ∼1 mL of DMEM, and then stored them at −80C.

We produced VSV-G pseudotyped viruses from the library cells as well. These viruses still contain the barcoded variants in their genomes, and can be used for PacBio sequencing or as controls for baseline variant frequencies during selections on the effects of mutations on Env entry into cells (Figure 2). We did this by plating 16 million library cells in a 15 cm plate and transfecting the plate 24 hours later using BioT according to the manufacturer’s recommendations. We used 7.5 ug of each lentivirus helper plasmid (Gag-Pol, Rev, and Tat) and a VSV-G expressing plasmid (four plasmids, 30 ug total DNA) for the transfection. 48 hours later we filtered the supernatant through a 0.45 µM SFCA filter and stored these viruses at −80C.

### PacBio sequencing of TRO.11 Env mutant libraries

We used long read PacBio sequencing to determine the TRO.11 Env mutant library composition and link the random nucleotide barcodes with the Env mutant in the same genomes, as described previously for the BF520 Env mutant libraries^18,26^. Briefly, we plated 1 million 293T cells per well in poly-L-lysine coated six well plates (Corning, Cat. No. 356515), and 24 hours later infected three wells of cells with ∼500,000 infectious units of +VSV-G pseudotyped library virus per well for each library. Three hours after infection, the media was replaced with fresh D10 to reduce cellular toxicity. Twelve hours after infection, we removed the media, washed the cells with PBS, and miniprepped each well to isolate the unintegrated lentivirus genomes^18,26,46^.

We amplified the barcoded TRO.11 Env sequences exactly as described previously for the BF520 Env mutant libraries^26^ using a two-step PCR protocol^18,26^. For the first round of PCR, the miniprepped lentivirus genomes were split into two short-cycle PCRs which each attached single nucleotide tags unique to each PCR to each end of the amplicons. The products from the first round PCRs were then pooled back together into a longer cycle second PCR. In this way, sequences of amplicons with mismatching nucleotide tags from the first round of PCR can be filtered out as having had strand exchange occur during the longer cycle PCR, and the total rate of strand exchange can be estimated. See Radford et al.^26^ for details on the primers and PCR cycling conditions.

### Selections on the effects of mutations on the function of HIV Env entry into cells

We measured the effects of mutations on Env mediated entry into cells exactly as described previously^26^ (Figure 4). Briefly, we infected 1 million TZM-bl cells per well in six well plates with ∼1 million infectious units of VSV-G or non-VSV-G pseudotyped mutant virus library, with 100 ug/mL of DEAE dextran. 12 hours after infection, we washed the cells with PBS, miniprepped the cells, and eluted into 30 µL of EB. We then prepared the eluents for barcode sequencing in the barcode sequencing preparation described below.

### Production and spike-in of neutralization standard VSV-G pseudotyped viruses

During selections on the effects of mutation on Env escape from antibodies, we spiked in a small amount of separately produced only-VSV-G pseudotyped virus carrying known barcodes to act as standards of infection, so that we could convert variant barcode frequencies to absolute neutralization values^18^. We produced these viruses exactly as described previously^18^. Briefly, 293T-rtTA cells were transduced at a low multiplicity of infection with VSV-G pseudotyped viruses carrying lentivirus genomes with known barcodes but no viral entry protein gene in their genomes. Transduced cells were sorted using flow cytometry, and then spike-in standard viruses were produced by transfecting these cells with lentiviral helper plasmids (Gag-Pol, Rev, and Tat) along with a separate plasmids expressing VSV-G.

### Selections on the effects of HIV Env mutations on escape from neutralization by antibodies

We measured the effects of mutations on Env escape from antibodies as described previously^18,26^. Briefly, we spiked-in the VSV-G pseudotyped neutralization standard into the genotype-phenotype linked virus libraries to be ∼1% of the total infectious units in the pool. For varying concentrations of antibody in the IC95-IC99.9 range and for a no-antibody control condition, we incubated 1 million infectious units of the combined library and standard virus pool with antibody (or no antibody for the control) for one hour. After the incubation, the volume of each incubation was raised to 2mL of D10 with 100 ug/mL DEAE dextran, and used to infect one well of TZM-bl cells at around one million cells per well in a six well dish. 12 hours after infection, the cells were washed with PBS, miniprepped, eluted into 30 µL of EB, and then prepared for barcode sequencing as described in the section below.

### Barcode sequencing preparation

After selections on virus libraries for the effects of mutations on cell entry or escape from antibodies, we prepared the samples for Illumina sequencing of barcodes exactly as described previously^18^. Briefly, we did an initial round of PCR to amplify the barcodes from the miniprep eluents using a forward primer that aligns to the Illumina Truseq read 1 sequence upstream of the barcode in the lentivirus genome and a reverse primer that anneals downstream of the barcode and overlaps with the Illumina Truseq read 2 sequence. We Ampure bead cleaned the products with a 1:3 product to beads ratio, and then followed this with a second round of PCR using a forward primer that anneals to the Illumina Truseq Read 1 sequence and has a P5 Illumina adapter overhang, and reverse primers from the PerkinElmer NextFlex DNA Barcode adaptor set that anneal to the Truseq Read 2 site and have the P7 Illumina adapter and an i7 sample index. The concentration of products from the second round of PCRs was quantified using Qubit, pooled at an even ratio, gel purified, Ampure bead cleaned at a 1:3 sample to beads ratio, and then sequenced using either P2 or P3 reagent kits on a NextSeq 2000. See Radford et al.^26^ for details on the primers and PCR cycling conditions.

### Single mutant neutralization curves

We performed traditional neutralization assays to validate the effects of mutations measured in our deep mutational scanning as described previously^26^. Briefly, rather than use ΔEnv HIV pseudoviruses in typical TZM-bl neutralization assays, for each validation mutant we co-transfected a plasmid that expresses Env with an escape escape mutation along with a ZsGreen and Luciferase expressing lentivirus backbone plasmid and lentiviral helper plasmids (Gag-Pol, Rev, and Tat). Upon infecting cells, these pseudoviruses express Luciferase from their lentivirus genomes, which we can quantify using the Bright-Glo Luciferase Assay System (Promega, E2610) to measure relative light units (RLUs). For titrations of these viruses and for the neutralization assays, we plated 25,000 TZM-bl cells per well in clear bottom, poly-L-lysine coated, black walled 96 well plates (Greiner, Cat. No. 655930). For titrations, we serially diluted each validation mutant virus, infected the cells, measured the RLUs per well, and then estimated the average RLU/µL for each validation mutant virus within a linear range based on its RLU dilution curve. For neutralization assays, we serially diluted the antibody, incubated each dilution with 200,00-400,000 RLUs of each validation mutant virus for one hour, added D10 with DEAE dextran to a final DEAE dextran concentration of 100ug/mL after the incubation, and infected the TZM-bl cells. 48 hours later, we measured the RLUs for each antibody dilution / validation mutant virus. See Radford et al.^26^ for more details.

## Computational Methods

### Data analysis pipeline overview and main outputs

For the main analysis of the deep mutational scanning data, we used *dms-vep-pipeline-3* (https://github.com/dms-vep/dms-vep-pipeline-3) version 3.16.3, a modular pipeline for analyzing viral entry protein deep mutational scanning data. See https://github.com/dms-vep/HIV_Envelope_TRO11_DMS_3BNC117_10-1074 and https://github.com/dms-vep/HIV_Envelope_BF520_DMS_3BNC117_10-1074 for all code and data used in the analysis.

See https://dms-vep.org/HIV_Envelope_TRO11_DMS_3BNC117_10-1074/ and https://dms-vep.org/HIV_Envelope_BF520_DMS_3BNC117_10-1074/ for HTML renderings of key analyses and interactive plots from the analysis.

See the table of key outputs and resources above for links to pre-filtered csv files containing the effects of mutations on entry into cells for TRO.11 Env and BF520 Env and for interactive structures shaded by the effects of mutations on entry into cells or antibody escape for TRO.11 Env and BF520 Env. We used *dms-viz*^47^ to make these interactive structures using PDB 9BER^43^ for all TRO11 Env mutation effects, PDB 6UDJ^48^ for BF520 Env cell entry and 10-1074 escape mutation effects, and PDB 5V8M^13^ for BF520 Env 3BNC117 escape mutation effects.

### Analysis of PacBio and Illumina sequencing

We analyzed the BF520 Env mutant libraries PacBio sequencing data in our prior study^26^. We analyzed the new TRO.11 Env mutant libraries PacBio sequencing data using the same data analysis pipeline. Briefly, we filtered out any CCSs with unexpected pairs of nucleotide tags from the PCRs before PacBio sequencing as likely strand-exchanged sequences, and estimated the total amount of strand exchange to be <1% of the total CCSs. We also filtered out any barcodes that had less than three CCSs or had minor fractions of substitutions or indels above 0.2, to remove barcodes that had either too few CCSs to confidently link them to an Env mutant or evidence of multiple Env mutants sharing that barcode. All barcoded variants for each replicate library that passed these filters were added to a barcode-variant lookup table. See https://dms-vep.org/HIV_Envelope_TRO11_DMS_3BNC117_10-1074/notebooks/build_pacbio_consensus.html for the analysis described in this section and see https://github.com/dms-vep/HIV_Envelope_TRO11_DMS_3BNC117_10-1074/blob/main/results/variants/codon_variants.csv for the barcode-variant lookup table.

We analyzed the Illumina barcode sequencing as described previously^26^. Briefly, we used the parser found at https://jbloomlab.github.io/dms_variants/dms_variants.illuminabarcodeparser.html/ to determine the counts of each variant in each selection condition. For each cell entry or antibody selection, we retained barcoded variants for downstream analyses if their pre-selection counts were above the thresholds specified in the configuration files found at https://github.com/dms-vep/HIV_Envelope_TRO11_DMS_3BNC117_10-1074/blob/main/data/func_effects_config.yml and https://github.com/dms-vep/HIV_Envelope_TRO11_DMS_3BNC117_10-1074/blob/main/data/antibody_escape_config.yml. This removes low frequency variants likely to have noisy scores due to random bottlenecking during experiments.

### Analysis of the effects of mutations on HIV Env function of entry into cells

We analyzed the effects of mutations on BF520 Env entry into cells in our prior study^26^, and here we analyzed the effects of TRO.11 Env mutations on entry into cells the same as previously^18,26^. Briefly, we calculated functional scores for each barcoded Env mutant relative to its parental wildtype Env, and then used global epistasis models^34,35^ to deconvolute the effects of single amino acid mutations from the barcoded variant functional scores. The deconvoluted single mutation effects on cell entry are the values reported in the figures of this manuscript and the csv files linked above.

### Analysis of the effects of mutations on HIV Env escape from neutralization by antibodies

We analyzed the effects of mutations on BF520 Env escape from neutralization by antibody 3BNC117 in our prior study^26^, and we analyzed the effects of mutations on escape for the rest of the Env / antibody pairs reported here the same as previously^18,26^. Briefly, we used the change in frequency of the spike-in standard viruses between the no-antibody and each antibody selection compared to the change in frequency of each barcoded variant to determine the fraction of each barcoded variant neutralized at each antibody concentration^18^. We used the software *polyclonal*^37^ to fit biophysical models of the effects of mutations on escape for each Env / antibody pair.

Under these models, each mutation has an individual effect on antibody escape, which are the escape values shown in the logo plots in Figures 4, 5, and 6, in the scatter plots in Figure 7, and in the heatmaps shown in Supplemental Figures 3, 4, 5, and 6. Using the models, these escape values can be transformed into the fraction of virus bearing a mutation neutralized at arbitrary antibody concentrations, which we used to calculate the deep mutational scanning-measured fold change IC50 values in Figures 5 and 6.

### Single mutant neutralization curve analysis

We analyzed the single mutant neutralization curves as described previously^26^. Briefly, we calculated the fraction infectivity of each single mutant virus at each antibody concentration by subtracting from each well’s RLUs the average background reading of RLUs from uninfected cells, and then dividing by the average RLUs from cells infected by the mutant virus that was incubated with media with no antibody. We used *neutcurve* (https://jbloomlab.github.io/neutcurve/) to fit neutralization curves with the fraction infectivities, plot the neutralization curves seen in Figures 5 and 6, and to calculate the neutralization assay measured IC50s see in Figures 5 and 6.

## Data availability

All data is available at links provided in the Methods section and in the table of key outputs and resources. The table of key outputs and resources includes csv outputs from the main analysis pre-filtered using suggested filters, as well as interactive structures of HIV Env shaded by deep mutational scanning measured mutation effects. See https://github.com/dms-vep/HIV_Envelope_TRO11_DMS_3BNC117_10-1074 and https://github.com/dms-vep/HIV_Envelope_BF520_DMS_3BNC117_10-1074 for the full analysis pipeline, inputs, and key results. See the Computational Methods section above for instructions on running the analysis. The raw sequencing data for this study can be found in the NCBI Sequence Read Archive under BioProject numbers PRJNA947170 and PRJNA1217050.

---

**Supplemental Figure 1:**
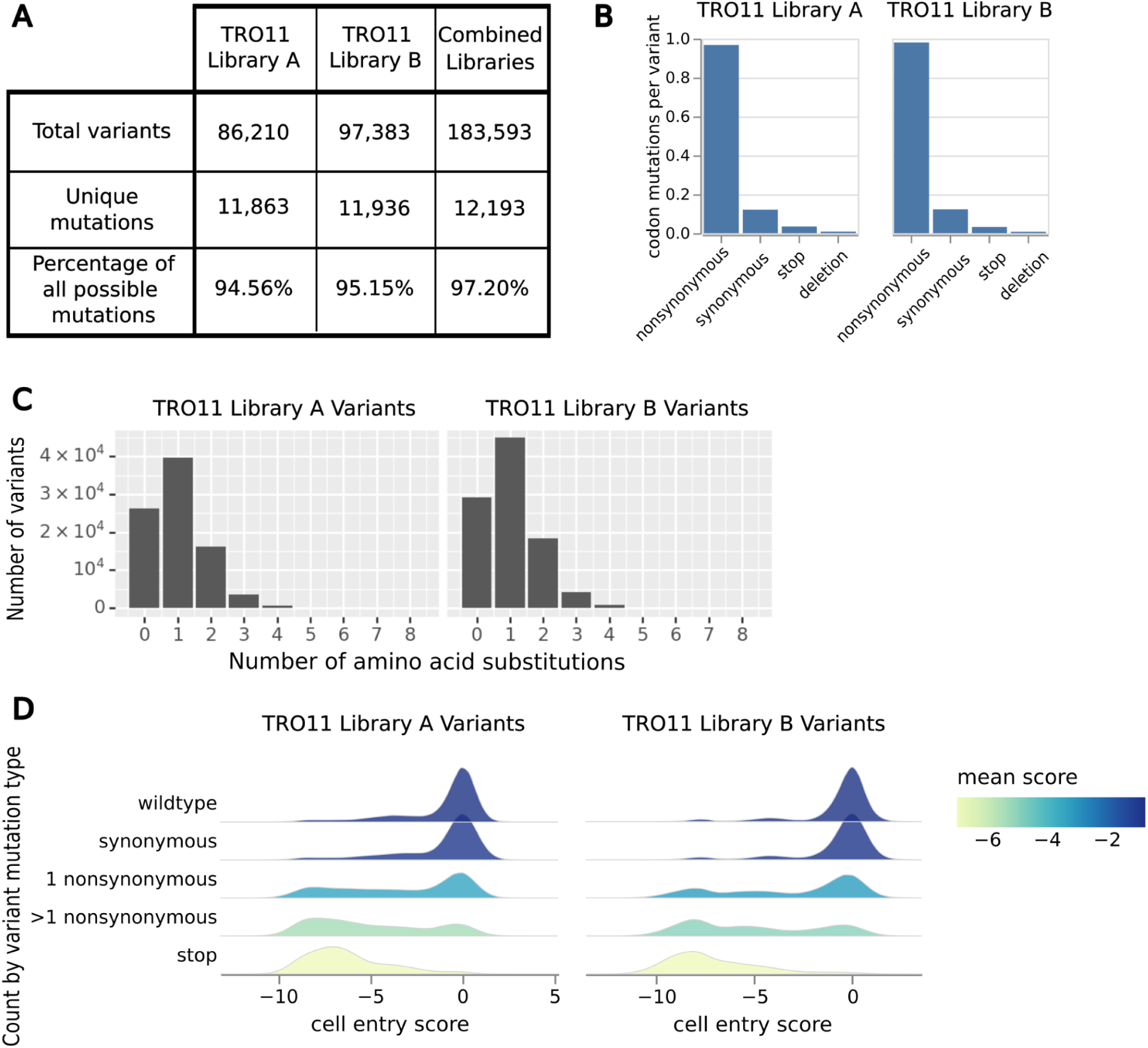
(A) Composition of variants and mutations in each individual TRO.11 Env mutant library and the combined TRO.11 libraries. (B) Average codon mutations per variant in each TRO.11 Env library, classified by type of codon mutation. (C) Distributions of the number of amino acid mutations per variant in the TRO.11 libraries. (D) Distributions of the cell entry scores of variants in each TRO.11 Env library, separated by types of codon mutations found in the mutants. Negative cell entry scores mean worse cell entry than the unmutated Env.

**Supplemental Figure 2:**
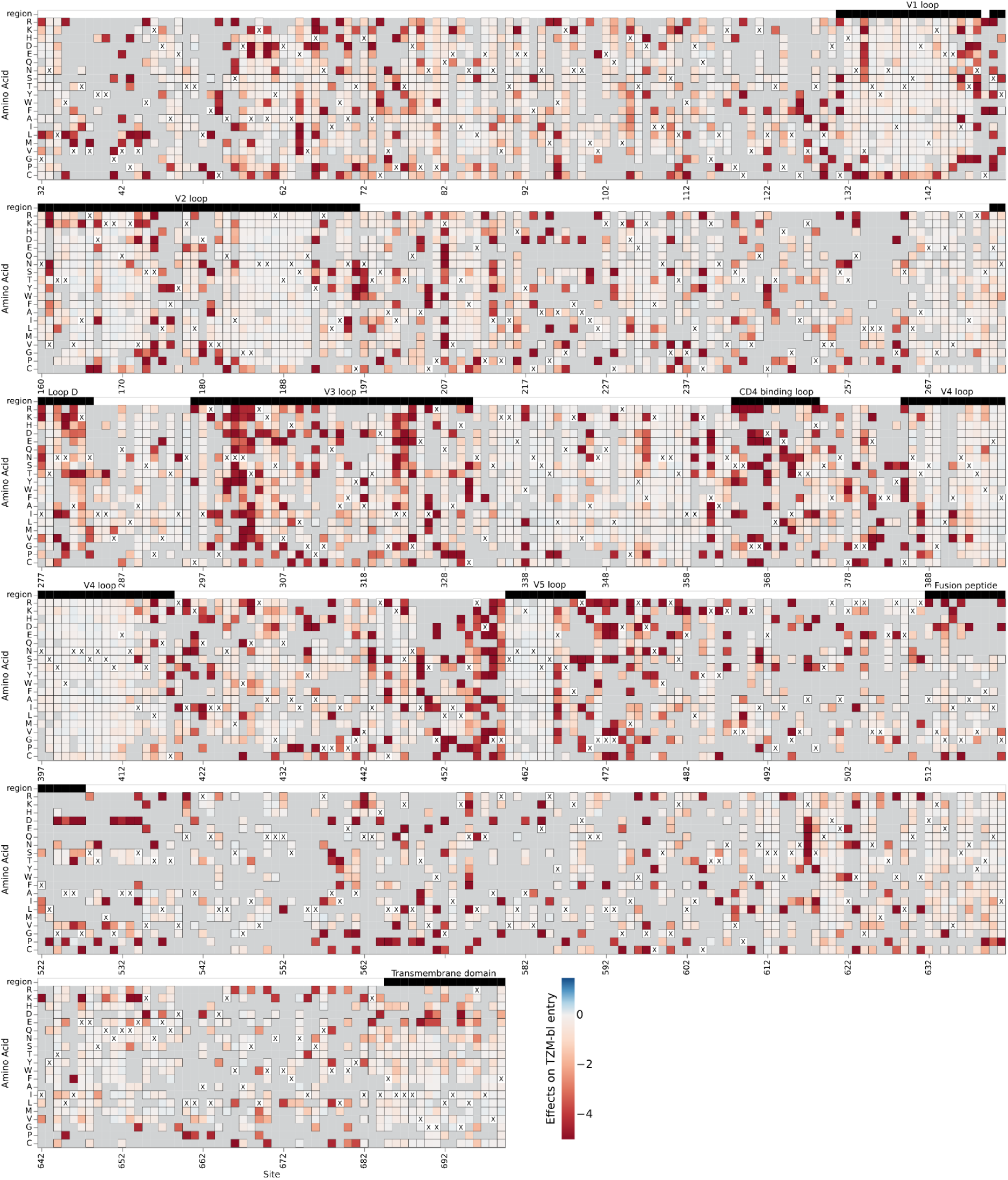
Effects of mutations on BF520 Env cell entry. For each mutation, the effect displayed in the heatmap is relative to unmutated BF520 Env. A value of zero (white) indicates no effect relative to unmutated BF520 Env, negative values (red) indicate impaired entry relative to unmutated BF520 Env, and positive values (blue) indicate improved entry relative to unmutated BF520 Env. Wildtype residues at each site are labeled with a black “X”. Notable regions of Env are denoted with labeled black bars above each row. Mutations with few observations in our experiments and therefore unconfident or no measurements are labeled with grey squares. See https://dms-vep.org/HIV_Envelope_BF520_DMS_3BNC117_10-1074/htmls/TZM-bl_entry_func_effects.html for an interactive version of this heatmap. BF520 Env deep mutational scanning measured effects of mutations on cell entry shown in this figure are previously published^26^.

**Supplemental Figure 3:**
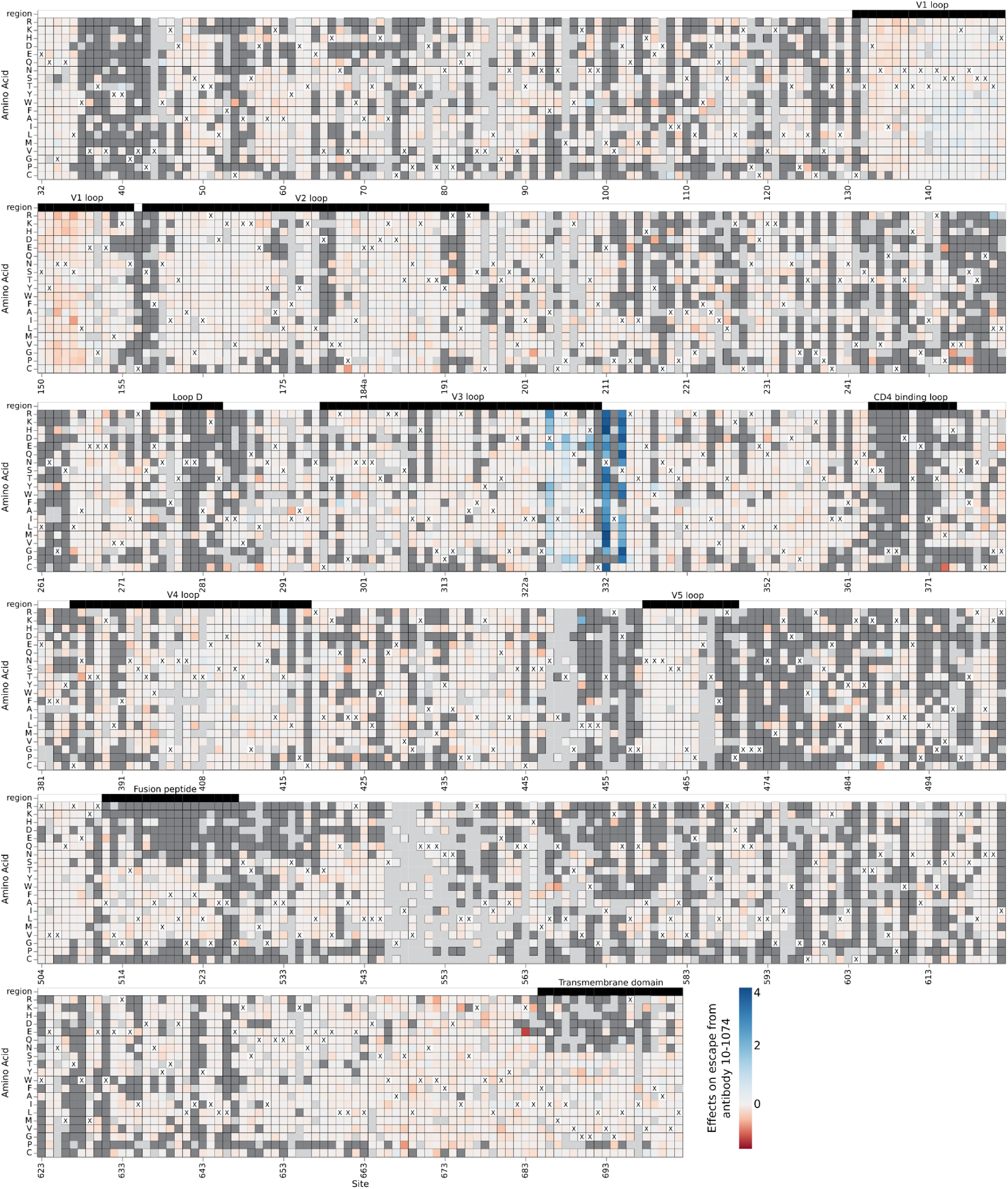
Effects of mutations on TRO.11 Env escape from antibody 10-1074. A mutation effect on escape of 10-1074 of zero (white) indicates no effect relative to unmutated TRO.11 Env, positive values (blue) indicate escape from neutralization by 10-1074 relative to unmutated TRO.11 Env, and negative values (red) indicate better neutralization by 10-1074 relative to unmutated TRO.11 Env. The parental amino-acid identity at each site in TRO.11 Env is labeled with a black “X”. Key regions of Env are denoted with labeled black bars above each row. Mutations that impair cell entry are colored dark gray, and mutations with effects that were not well measured in our experiments are colored light gray. See https://dms-vep.org/HIV_Envelope_TRO11_DMS_3BNC117_10-1074/htmls/10-1074_mut_effect.html for an interactive version of this heatmap.

**Supplemental Figure 4:**
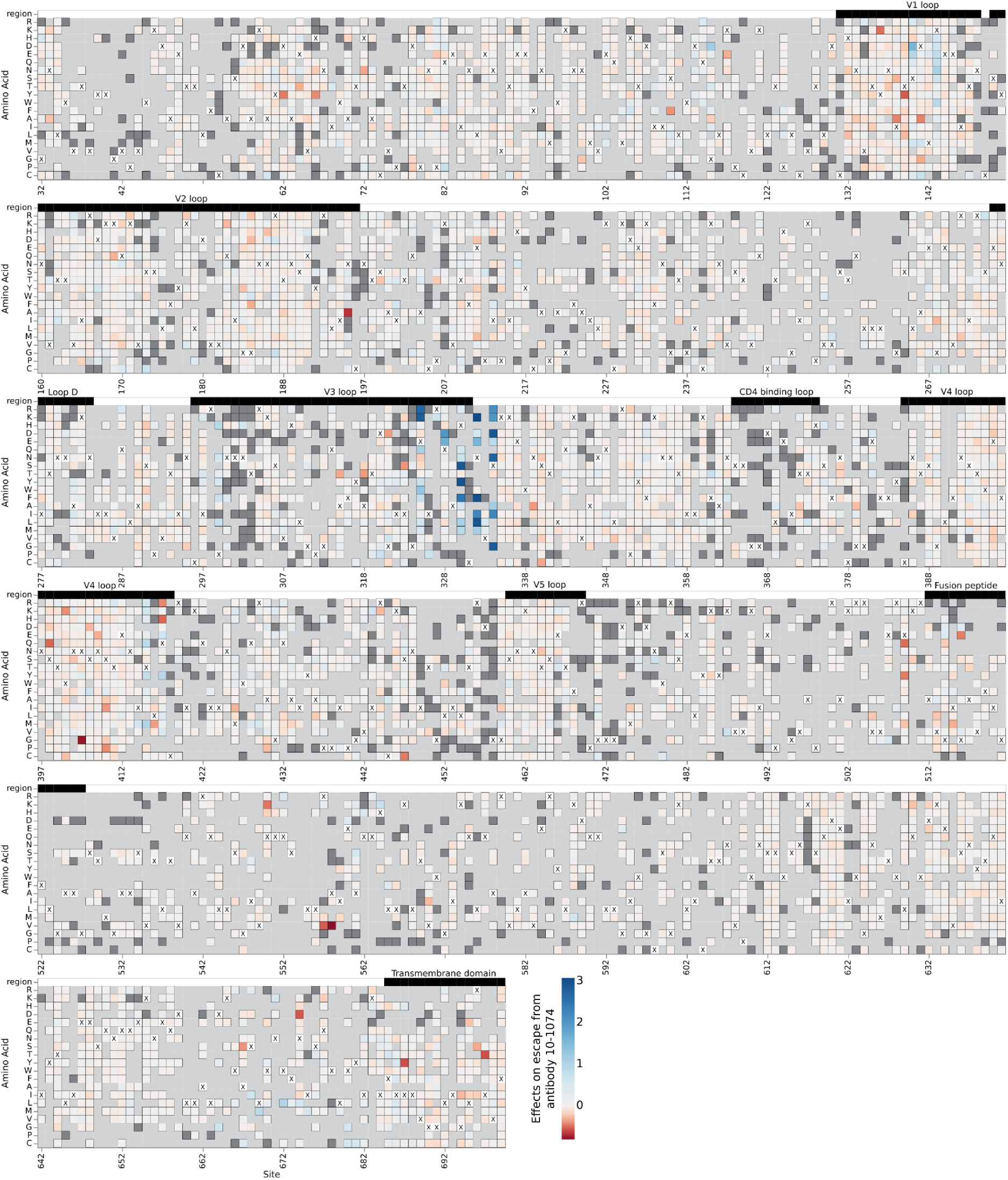
Effects of mutations on BF520 Env escape from antibody 10-1074. A mutation effect on escape of 10-1074 of zero (white) indicates no effect relative to unmutated BF520 Env, positive values (blue) indicate escape from neutralization by 10-1074 relative to unmutated BF520 Env, and negative values (red) indicate higher neutralization by 10-1074 relative to unmutated BF520 Env. The parental amino-acid identity at each site in BF520 Env is labeled with a black “X”. Key regions of Env are denoted with labeled black bars above each row. Mutations that impair cell entry are colored dark gray, and mutations with effects that were not well measured in our experiments are colored light gray. See https://dms-vep.org/HIV_Envelope_BF520_DMS_3BNC117_10-1074/htmls/10-1074_mut_effect.html for an interactive version of this heatmap.

**Supplemental Figure 5:**
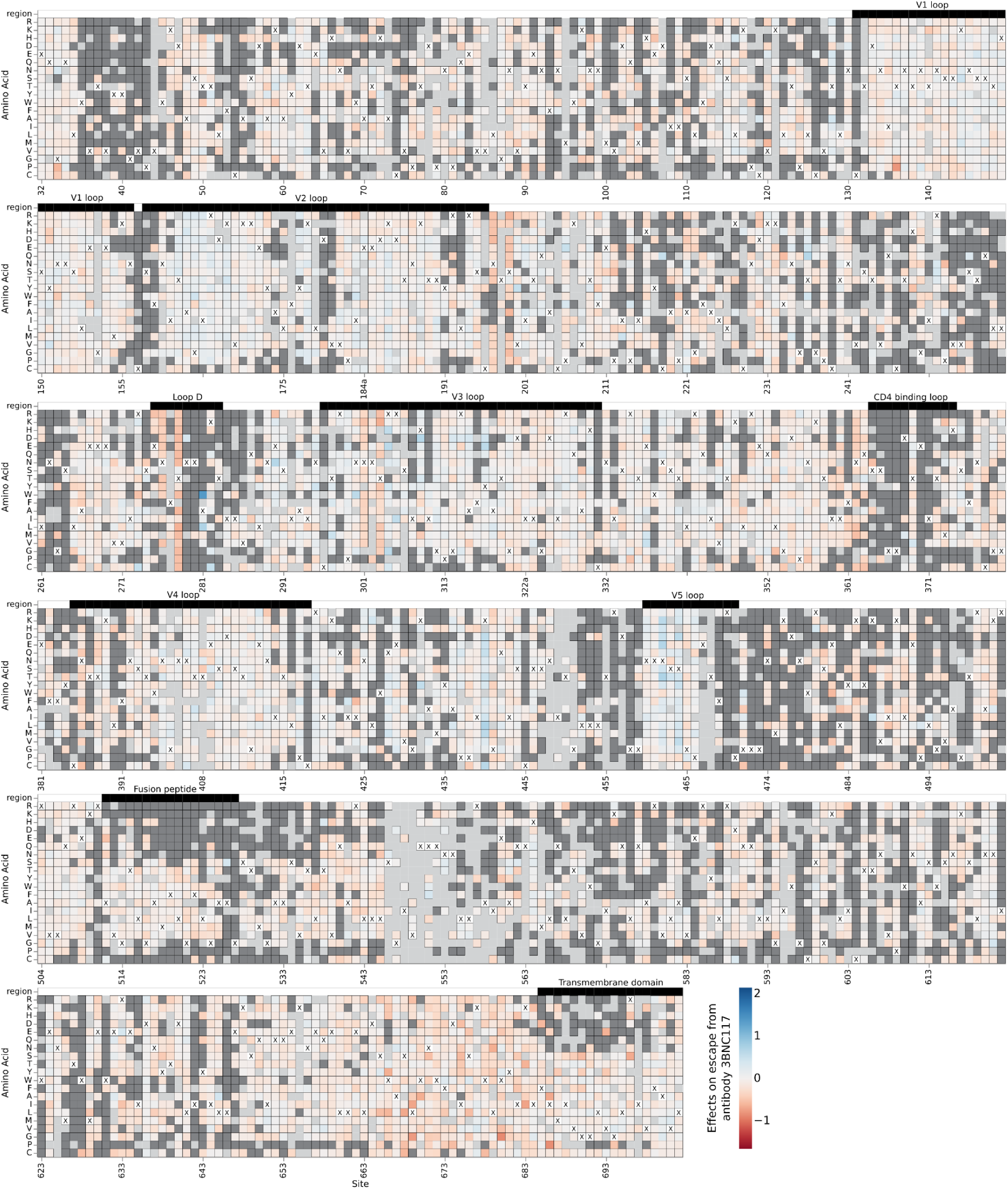
Effects of mutations on TRO.11 Env escape from antibody 3BNC117. A mutation effect on escape of 3BNC117 of zero (white) indicates no effect relative to unmutated TRO.11 Env, positive values (blue) indicate escape from neutralization by 3BNC117 relative to unmutated TRO.11 Env, and negative values (red) indicate higher neutralization by 3BNC117 relative to unmutated TRO.11 Env. The parental amino-acid identity at each site in TRO.11 Env is labeled with a black “X”. Key regions of Env are denoted with labeled black bars above each row. Mutations that impair cell entry are colored dark gray, and mutations with effects that were not well measured in our experiments are colored light gray. See https://dms-vep.org/HIV_Envelope_TRO11_DMS_3BNC117_10-1074/htmls/3BNC117_mut_effect.html for an interactive version of this heatmap.

**Supplemental Figure 6:**
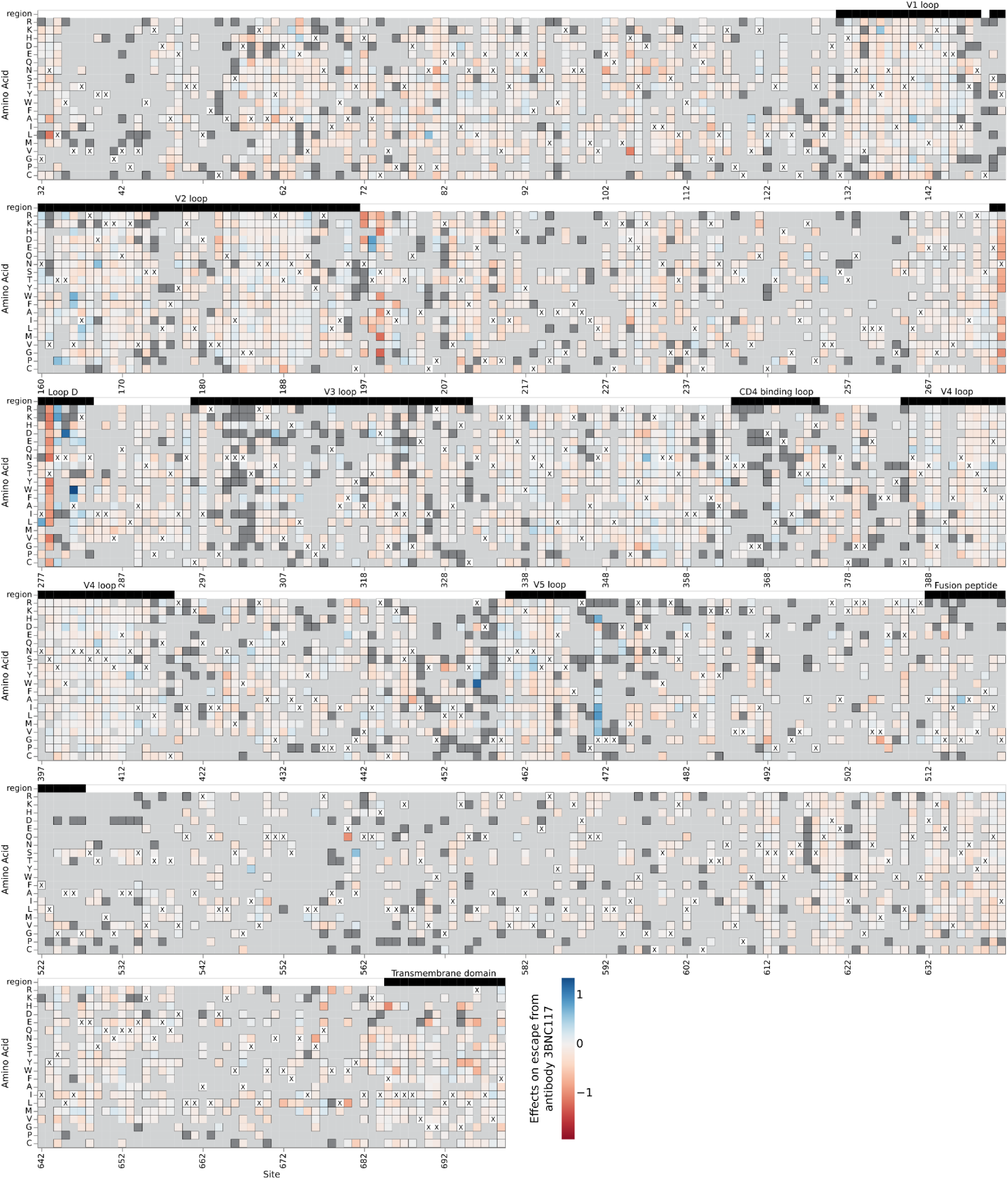
Effects of mutations on BF520 Env escape from antibody 3BNC117. A mutation effect on escape of 3BNC117 of zero (white) indicates no effect relative to unmutated BF520 Env, positive values (blue) indicate escape from neutralization by 3BNC117 relative to unmutated BF520 Env, and negative values (red) indicate higher neutralization by 3BNC117 relative to unmutated BF520 Env. The parental amino-acid identity at each site in BF520 Env is labeled with a black “X”. Key regions of Env are denoted with labeled black bars above each row. Mutations that impair cell entry are colored dark gray, and mutations with effects that were not well measured in our experiments are colored light gray. See https://dms-vep.org/HIV_Envelope_BF520_DMS_3BNC117_10-1074/htmls/3BNC117_mut_effect.html for an interactive version of this heatmap. BF520 Env deep mutational scanning measured effects of mutations on escape from antibody 3BNC117 shown in this figure are previously published^26^.

